# Polycomb shapes active chromatin and promoter bivalency during ovarian reserve formation and activation

**DOI:** 10.64898/2026.04.22.720278

**Authors:** Mengwen Hu, Yasuhisa Munakata, Yu-Han Yeh, Neil Hunter, Richard M. Schultz, Satoshi H. Namekawa

## Abstract

The ovarian reserve, a finite pool of long-lived non-growing oocytes established at birth, determines female reproductive lifespan, yet how these oocytes establish long-term quiescence while retaining the capacity for future growth and embryogenesis remains poorly understood. Here, we define a regulatory logic by which Polycomb repressive complexes shape stage-specific active chromatin remodeling during ovarian reserve formation and early oocyte growth. During ovarian reserve formation, H3K27ac, an active promoter- and enhancer-associated mark, undergoes extensive genome-wide redistribution. A key feature of this transition is CpG island promoter remodeling, in which many loci lose H3K27ac while gaining PRC1-dependent H2AK119ub, a repressive mark. This early reprogramming is followed during oocyte growth by acquisition of PRC2-dependent H3K27me3, de novo establishment of bivalent promoters, and protection of promoter regions from de novo DNA methylation. Oocyte growth is also accompanied by broad gains in both H3K27ac and H3K4me3, an active promoter-associated mark. Analyses of PRC1- and PRC2-deficient oocytes reveal unequal Polycomb contributions: PRC2 broadly constrains H3K27ac, whereas PRC1 more selectively shapes genome-wide H3K27ac redistribution and restricts H3K4me3 accumulation at bivalent promoters. Together, these findings identify staged active chromatin remodeling as an integral feature of perinatal oocyte development and reveal that Polycomb shapes chromatin state transitions as oocytes enter quiescence and become poised for future growth.

**One-Sentence Summary:** Polycomb repressive complexes shape stage-specific active chromatin remodeling to establish quiescence and future promoter states during ovarian reserve formation and early oocyte growth.

## Introduction

The ovarian reserve is a finite pool of oocytes that sustains female fertility across reproductive lifespan (*1–3*). Established around birth, this pool consists of long-lived non-growing oocytes (NGOs) enclosed within primordial follicles (*3–8*). Although often viewed as a dormant reservoir, the ovarian reserve represents a more demanding developmental state: oocytes must acquire long-term quiescence while preserving the capacity to resume growth, complete oogenesis, and ultimately support fertilization and early embryogenesis. A central question in female germline biology is how oocytes establish an epigenomic state that sustains quiescence while preserving competence for future growth and embryogenesis.

In mice, formation of the ovarian reserve is followed by recruitment of a subset of primordial follicles with initiation of oocyte growth, giving rise to growing oocytes (GOs) that later mature into fertilizable eggs (*9, 10*). These transitions are accompanied by extensive transcriptional reprogramming (*11–14*), indicating that progression from embryonic oocytes to NGOs and then to GOs is governed by a coordinated developmental program rather than by simple entry into and release from quiescence. Defining chromatin mechanisms that operate across these transitions is therefore essential for understanding how the ovarian reserve is established, maintained, and linked to subsequent oocyte growth.

Following genome-wide DNA demethylation in primordial germ cells, the oocyte genome remains globally hypomethylated until postnatal growth, when de novo DNA methylation and specialized chromatin landscapes are progressively established (*15–18*). Full-grown oocytes (FGOs) display unusual epigenomic features, including broad H3K4me3 domains (*19, 20*), a histone modification typically enriched at active promoters, noncanonical H3K27ac patterns (*18, 21*), a mark associated with active promoters and enhancers, and widespread H3K27me3 (*22*), a repressive histone modification deposited by PRC2 that persists into early embryogenesis and contributes to DNA methylation–independent imprinting (*23–25*). The developmental origins of these distinctive chromatin states, however, remain poorly understood, particularly during the perinatal period, when oocytes transition from embryonic meiotic prophase I into the ovarian reserve and subsequently into the growth phase (*26*).

Polycomb repressive complexes are central regulators of developmental chromatin states (*27–29*). In the female germline, PRC1-mediated H2AK119ub-dependent repression of the meiotic prophase I (MPI) program underlies proper ovarian reserve formation and is linked to later establishment of H3K27me3 during oocyte growth (*14, 30*). Together, these studies established Polycomb-mediated chromatin programming as a key feature of perinatal oogenesis but left unresolved whether Polycomb acts only to impose repressive chromatin or also shapes chromatin features associated with future activation competence. This question is especially relevant in the context of active chromatin, because H3K4me3 and H3K27ac are generally associated with transcriptional competence, whereas Polycomb is classically linked to repressive chromatin marked by H2AK119ub and H3K27me3.

Here, by analyzing PRC1- and PRC2-deficient oocytes, we define the molecular logic by which Polycomb repressive complexes shape stage-specific remodeling of active chromatin during ovarian reserve formation and oocyte growth. Our findings identify promoter remodeling and subsequent bivalent promoter priming as integral features of perinatal oocyte development and suggest that Polycomb shapes chromatin state transitions as oocytes enter quiescence and become poised for future growth.

## Results

### Dynamic active chromatin remodeling during perinatal oogenesis

In mice, perinatal oocytes undergo two key developmental transitions: the perinatal oocyte transition (POT), during which oocytes exit MPI and establish the ovarian reserve (*14*), and the primordial-to-primary follicle transition (PPT), during which a subset of primordial follicles initiates growth (*13*). To define active chromatin dynamics across the POT and PPT, we profiled H3K4me3 and H3K27ac at four key stages of mouse perinatal oogenesis: E18.5 oocytes in MPI (E18O), oocytes transitioning to dictyate arrest at postnatal day 1 (P1O), NGOs within primordial follicles at P6 (NGO), and GOs in primary follicles at P7 (GO) (Fig. 1A). To quantify stage-specific changes, we performed low-input cleavage under targets and tagmentation (CUT&Tag) (*31, 32*) with spike-in controls normalized to cell number, enabling quantitative comparison across stages (fig. S1A). After confirming dataset quality and reproducibility (fig. S1B), biological replicates were pooled for downstream analyses.

**Fig. 1.**
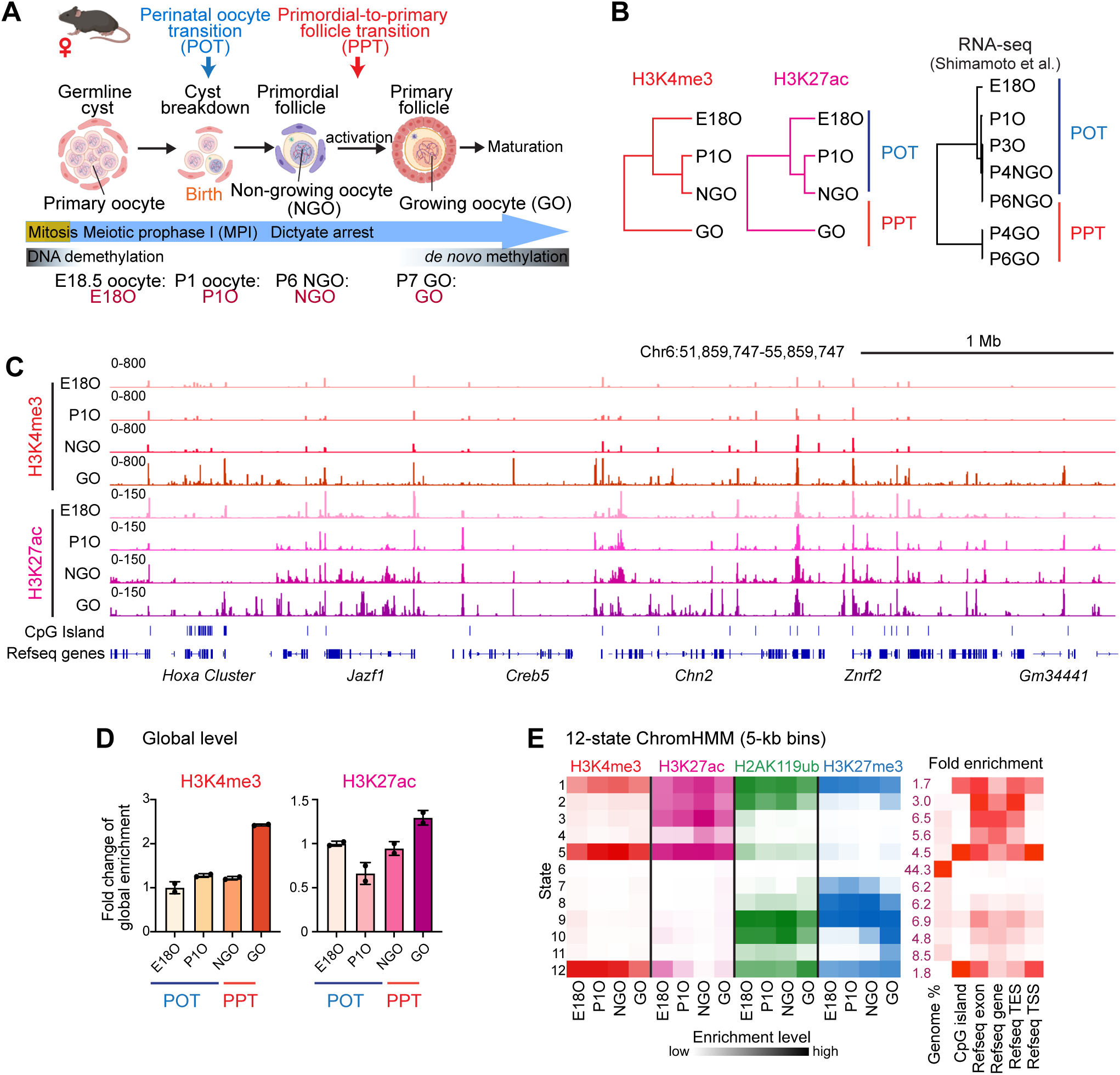
Global remodeling of H3K27ac and H3K4me3 during POT and PPT. (A) Schematic of mouse perinatal oogenesis (created with BioRender.com) also depicting transition from mitosis to meiosis and changes in DNA methylation levels. NGO, non-growing oocyte; GO, growing oocyte. The stages analyzed in this study are indicated at the bottom. E18O represents oocytes in meiotic prophase I; P1O and P3O represent oocytes transitioning to dictyate arrest; P4 and P6 small oocytes represent NGO residing in primordial follicles; and P4 and P6 large oocytes represent GO in primary follicles after initiation of oocyte growth. (B) Hierarchical clustering of global H3K4me3 and H3K27ac in 10-kb bins and hierarchical clustering of gene expression across all stages during perinatal oogenesis in wild-type. (C) Track views of H3K4me3 and H3K27ac landscapes during perinatal oogenesis. The data ranges represent spike-in scaled RPKM (sRPKM) values. (D) Bar chart showing the fold change of global levels of H3K4me3 and H3K27ac as determined by quantitative CUT&Tag in perinatal oocytes (n = 2 biological replicates, indicated by dots). (E) Heatmap illustrating chromatin state dynamics in perinatal oocytes using ChromHMM analysis. The intensity of colors indicates the enrichment level for each modification belonging to the given state at each stage. Genome distributions of all 12 states were shown on the right. The intensity of colors indicates the fold enrichment level of the given state. The number in the first column represents the genome coverage percentage of each state. H2AK119ub and H3K27me3 CUT&Tag data were reanalyzed from Hu et al. (*30*).

Hierarchical clustering revealed two major epigenomic transitions in active chromatin corresponding to the POT and PPT (Fig. 1B), accompanied by dynamic changes in the oocyte transcriptome (Fig. 1B, right) and in expression of enzymes that regulate active histone marks (fig. S1C). During POT, H3K4me3 levels remained relatively constant (Fig. 1, C and D), whereas H3K27ac underwent extensive redistribution from E18O to NGO (Fig. 1, C and D). By contrast, during PPT, both H3K27ac and H3K4me3 increased more broadly, consistent with broad transcriptional activation during early oocyte growth (Fig. 1, C and D).

Peak-based analyses corroborated these stage-specific changes. H3K4me3 peaks displayed relatively limited changes during POT but broadened and accumulated more extensively during PPT (fig. S2, A to D). In contrast, H3K27ac peaks showed more prominent changes in peak number, genomic distribution, and differential enrichment during POT (fig. S2, E to H), indicating dynamic H3K27ac remodeling during ovarian reserve establishment. Notably, gains of H3K27ac peaks in NGOs were observed in intergenic regions and gene bodies, suggesting acquisition of distal regulatory elements during POT (fig. S2, E and H).

These dynamic epigenomic transitions were further supported by ChromHMM analysis (*33*), which integrated both active and Polycomb-mediated repressive chromatin features across these developmental stages (Fig. 1E). By classifying genomic regions into 12 chromatin states, we identified dynamic, region-specific epigenomic changes during perinatal oogenesis. Notably, states 1, 5, and 12 were characterized by high levels of H3K4me3 and H3K27ac and enrichment for CpG islands (CGIs), consistent with promoter-like regions. States 1 and 12 were additionally enriched for the repressive marks H2AK119ub and H3K27me3, consistent with putative bivalent promoters. ChromHMM further confirmed genome-wide redistribution of H3K27ac during POT, particularly the extensive loss of state 12 signal at CGI-enriched promoters and gains of states 3 and 4 outside putative promoters.

Together, these data indicate that ovarian reserve formation is marked by H3K27ac redistribution during POT, followed by broader increases in both H3K27ac and H3K4me3 during subsequent oocyte growth. Furthermore, the abundance of active marks in NGOs suggests that these cells are not simply dormant but retain active regulatory elements despite remaining quiescent.

### Coordinated chromatin and transcriptional transitions during the POT and PPT

In mammals, a large group of promoters contains CGIs that are generally constitutively unmethylated and enriched for PRC1 and PRC2, as well as the H2AK119ub and H3K27me3 marks they mediate, respectively (*34, 35*). Following our observation of H3K27ac loss at CGI-enriched promoter regions, we compared histone marks at 14,040 CGI-associated genes (CGI genes, including 11,399 active genes and 2,641 inactive genes in perinatal oocytes) and 39,755 genes without CGIs (non-CGI genes, including 8,694 active genes and 31,061 inactive genes in perinatal oocytes), and found extensive promoter reprogramming at CGI genes (Fig. 2A). Overall, promoters of active genes were associated with high levels of active marks, whereas inactive genes were enriched for repressive marks. Strikingly, a sharp loss of H3K27ac was observed at ∼62% (n = 8,615) of CGI gene TSSs during POT regardless of gene expression status (Fig. 2B), whereas H3K4me3 levels remained relatively constant (Fig. 2, A and B). These changes were unexpected given the extensive genome-wide overlap between H3K27ac and H3K4me3 peaks (fig. S2I). Consistent with our recent report (*30*), H2AK119ub increased extensively and peaked in NGOs, whereas H3K27me3 enrichment followed this pattern and peaked in GOs (Fig. 2A). In particular, gain of H3K27me3 at CGI promoters in GOs appeared to coincide with substantial de novo bivalent promoter formation, defined by the coexistence of H3K4me3 and H3K27me3. In addition, H3K4me3 and H3K27ac increased during PPT, consistent with global gene activation in GOs.

**Fig. 2.**
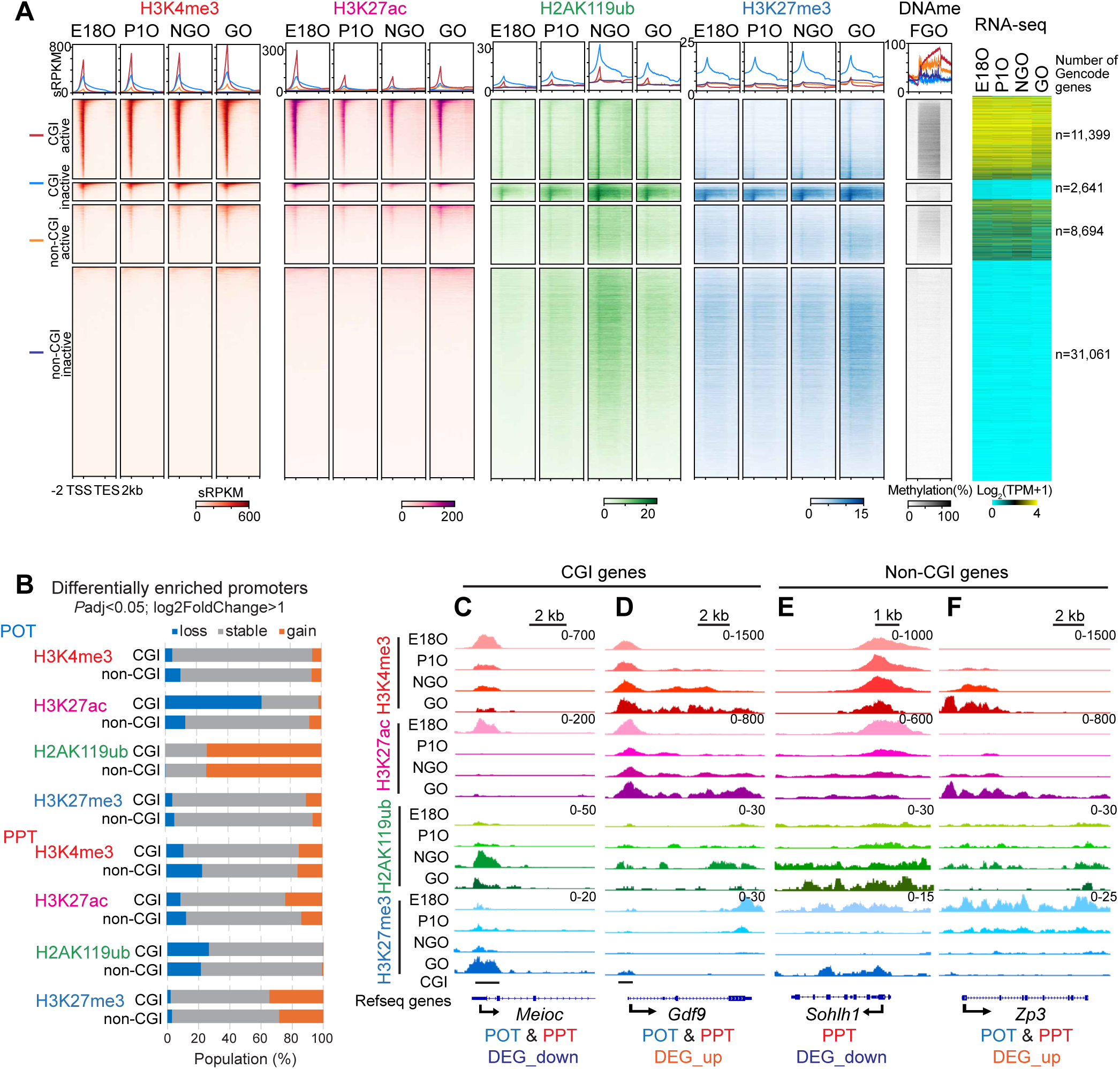
Coordinated chromatin and transcriptional transitions during POT and PPT. (A) Average tag density plots and heatmaps showing H3K4me3, H3K27ac,H2AK119ub, and H3K27me3 dynamics at gene body regions (TSS -TES ± 2 kb) of CGI genes and non-CGI genes in perinatal oocytes. The color keys represent signal intensity, and the numbers represent spike-in scaled RPKM (sRPKM) values. Right: average tag density plot and heatmap showing DNA methylation levels in full-grown oocytes (FGO) of corresponding CGI and non-CGI gene groups. The color keys represent signal intensity, and the numbers represent DNA methylation percentage. Heatmap showing RNA expression of corresponding genes in perinatal oocytes. The color keys represent signal intensity, log2-transformed TPM values. (B) Bar charts showing percentages of differentially enriched promoters of CGI and non-CGI genes for H3K4me3, H3K27ac, H2AK119ub, and H3K27me3 during POT and PPT. (C-F) Track views of the representative POT or PPT DEGs in CGI gene group (*Meioc* and *Gdf9*) and non-CGI gene group (*Sohlh1* and *Zp3*) gene loci showing differential H2AK119ub, H3K27me3, H3K4me3, and H3K27ac occupancy during perinatal oogenesis. Data ranges in the upper right represent sRPKM values from combined replicates.

To examine the relationship between chromatin state and gene expression during POT and PPT, we analyzed differentially expressed genes (DEGs) across these transitions using publicly available data (*11*) (fig. S3A). Genome-wide analysis revealed that global loss of H3K27ac and gain of H2AK119ub at promoters during POT were frequently associated with both up- and down-regulated genes (fig. S3, B and C). *Meioc* is a representative down-regulated CGI gene and a critical regulator of MPI (*36*) that is expressed in MPI and repressed during POT and PPT (fig. S3A). At the *Meioc* promoter, loss of H3K27ac and gain of H2AK119ub were observed during POT, followed by gain of H3K27me3 in GOs (Fig. 2C). *Gdf9* is a representative up-regulated CGI gene and an essential regulator of folliculogenesis (*37*) whose expression increases during POT and PPT (fig. S3A). During POT, loss of H3K27ac and gain of H2AK119ub were observed at the *Gdf9* promoter, whereas H3K4me3 and H3K27ac increased across the gene body as expression increased (Fig. 2D). Together, these findings indicate that promoter reprogramming occurs at both up- and down-regulated genes, suggesting that a relatively repressive promoter chromatin state is established after POT and may contribute to the long-term stability of the ovarian reserve.

Notably, CGI promoter reprogramming precedes DNA methylation patterning associated with transcription-dependent de novo DNA methylation in GOs (*38*). Reanalysis of previous whole-genome bisulfite sequencing data from FGOs (*15*) showed that, at CGI genes, DNA methylation increased globally across gene bodies while TSSs remained unmethylated (Fig. 2A). Thus, promoter reprogramming at CGI genes during POT, characterized by loss of H3K27ac and gain of H2AK119ub, together with subsequent gain of H3K27me3 during PPT, may contribute to protection of TSSs from de novo DNA methylation in GOs and to later DNA methylation patterning in FGOs.

In contrast to CGI genes, non-CGI genes displayed distinct gene-body-associated chromatin regulation. Within gene bodies of non-CGI genes, H3K4me3 and H3K27ac signals were generally sparse, whereas H2AK119ub increased globally in NGOs, followed by gain of H3K27me3 in GOs (Fig. 2A). *Sohlh1* is a representative non-CGI gene required for oogenesis (*39*) that is expressed in NGOs and down-regulated during PPT. Its down-regulation in GOs was associated with decreased H3K27ac and H3K4me3 and increased H2AK119ub and H3K27me3 (Fig. 2E). Another non-CGI gene, *Zp3*, encoding Zona Pellucida 3 (ZP3), which is essential for fertility (*40, 41*), is expressed in both NGOs and GOs. Similar to *Sohlh1*, H2AK119ub covered the gene body when *Zp3* was expressed in NGOs (Fig. 2F). However, *Zp3* induction in GOs was associated with reduced H2AK119ub and H3K27me3 and increased H3K4me3 and H3K27ac. Thus, CGI and non-CGI genes undergo distinct modes of chromatin reprogramming during perinatal oogenesis, with CGI genes showing promoter-centered remodeling and non-CGI genes showing gene-body-associated changes.

### H3K27ac reprogramming during POT is accompanied by gain of H2AK119ub

H3K27ac reprogramming during POT was not unidirectional, but instead involved selective loss, gain, and maintenance of H3K27ac across the genome. Differential peak analysis between E18O and NGOs identified three major classes of H3K27ac peaks (Fig. 3A): 17,481 peaks that lost H3K27ac during POT (“POT loss” peaks), 33,667 peaks that gained H3K27ac (“POT gain” peaks), and 62,127 peaks that remained relatively stable (“POT stable” peaks). These three classes included both proximal and distal elements, with proximal peaks notably enriched among “POT loss” peaks (Fig. 3B). “POT loss” peaks were enriched for YY1 and MAFK/MAFB motifs, whereas “POT gain” peaks were enriched for motifs recognized by TCF3 (E2A), PTF1A, and TCF12 (HEB) (Fig. 3C). YY1 functions as a Polycomb-associated factor (*42*) and is essential for follicle expansion (*43*), whereas TCF3 and TCF12 are critical regulators of postnatal oogenesis and folliculogenesis (*18*). Thus, H3K27ac reprogramming during POT is associated with developmental rewiring of gene regulatory elements.

**Fig. 3.**
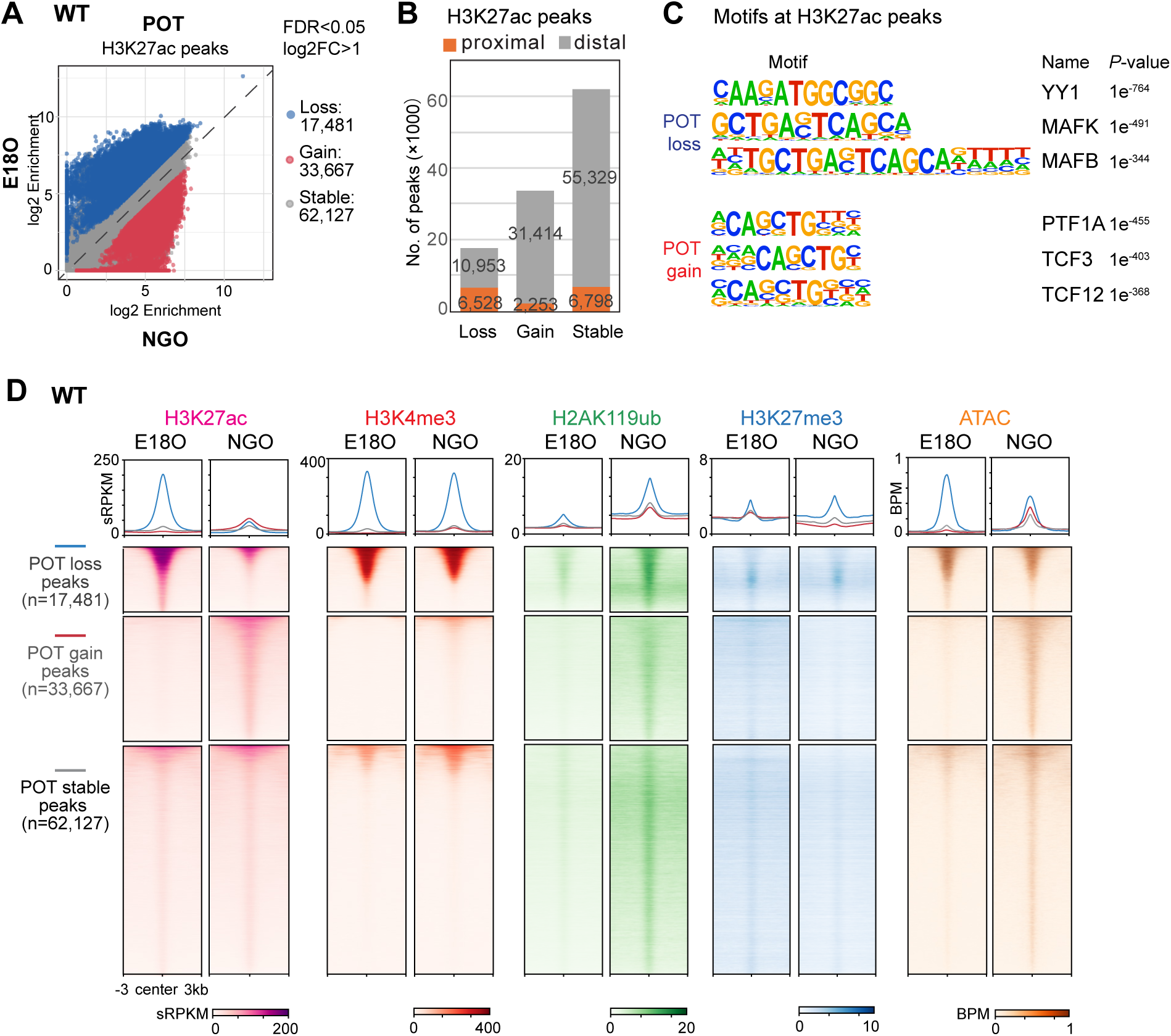
Global H3K27ac reprogramming during ovarian reserve formation. (A) Scatter plot showing the comparison of H3K27ac enrichment at the peak regions between E18O and NGO in WT. Colored dots represent peaks with differential H3K27me3 enrichment (FDR < 0.05, log2FoldChange>1). (B) Bar chart showing distribution of H3K27ac peaks at the proximal and distal regions. (C) Motif analysis of differentially enriched H3K27ac peaks during POT (*P* value, hypergeometric test with Bonferroni correction, one-sided, from HOMER; Methods). (D) Average tag density plots and heatmaps showing histone modification and chromatin accessibility changes at differential H3K27ac peak regions (peak center ± 3 kb) in E18O and NGO of WT. The color keys represent signal intensity, and the numbers represent spike-in scaled RPKM (sRPKM) values. ATAC-seq data were reanalyzed from Munakata et al. (*89*).

Notably, POT loss peaks gained H2AK119ub but lost chromatin accessibility during POT (Fig. 3D). At these peaks, H3K4me3 intensity remained relatively high, likely reflecting the enrichment of proximal elements (fig. S4A). By contrast, “POT gain” and “POT stable” peaks showed only modest gains in H2AK119ub and accessibility in NGOs. These patterns raised the possibility that Polycomb contributes to H3K27ac reprogramming during POT. Consistent with this idea, PRC2-mediated H3K27me3 is mutually exclusive with H3K27ac because the two modifications occupy the same residue, whereas PRC1 may influence H3K27ac through interactions with histone acetyltransferase (*44*) or deacetylase activities (*45*).

### PRC1 regulates H3K27ac reprogramming during ovarian reserve formation

To determine how Polycomb complexes contribute to H3K27ac reprogramming, we used two germ cell-specific conditional knockout mouse models that disrupt PRC1 or PRC2 function. Our previously established PRC1 loss-of-function model (PRC1cKO) and PRC2 loss-of-function model (PRC2cKO) (*14, 30, 46, 47*) use *Ddx4*-Cre, which is expressed in germ cells from E15 (*48*), to inactivate RNF2, the catalytic E3 ubiquitin ligase of PRC1, in a genetic background lacking the partially redundant paralog RING1 (RING1A) (*49, 50*), or EED, an essential component of PRC2 (*51*), respectively (Fig. 4, A and B). As reported previously, these models result in depletion of H2AK119ub or H3K27me3 in NGOs, respectively (*14, 30*). We therefore examined H3K27ac profiles in PRC1- and PRC2-deficient NGOs (fig. S5).

**Fig. 4.**
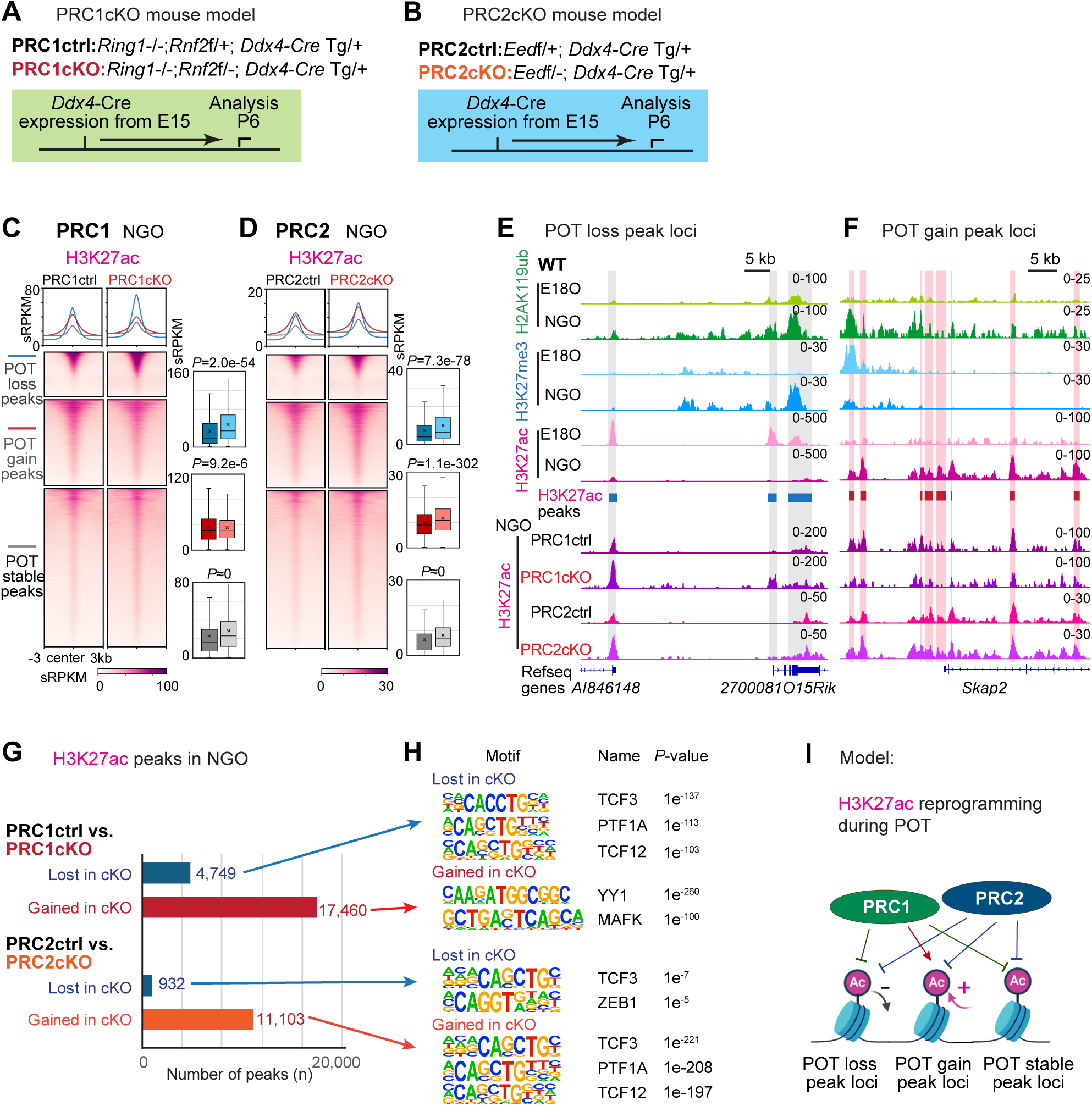
PRC1 regulates H3K27ac reprogramming during ovarian reserve formation. (A and B) Schematic of mouse models and experiments. (C and D) Average tag density plots and heatmaps showing H3K27ac changes at differential H3K27ac peak regions (peak center ± 3 kb) in NGO of PRC1ctrl&cKO (C), NGO of PRC2ctrl&cKO (D), respectively. Right: Box and whisker plots showing quantifications of H3K27ac enrichment within peaks (sRPKM values) of indicated peak groups in PRC1ctrl&cKO, PRC2ctrl&cKO, respectively. Boxes show the 25^th^ and 75^th^ percentile with the median, and whiskers indicate 1.5 times the interquartile range. *P* values of pairwise comparisons (two-sided unpaired Student’s t-test) are given. (E and F) Track views of the representative POT loss (E) or POT gain (F) H3K27ac peak regions showing H2AK119ub, H3K27me3, and H3K27ac occupancy during POT in WT, and H3K27ac changes in PRC1ctrl&cKO, PRC2ctrl&cKO NGO. Data ranges represent sRPKM values from combined replicates. (G and H) Bar chart showing numbers (G) and motif analysis (H) of differentially enriched H3K27ac peaks in NGO of PRC1ctrl&cKO, PRC2ctrl&cKO. (I) A schematic model showing the regulation of H3K27ac reprogramming during POT by Polycomb.

Loss of PRC1 and PRC2 affected H3K27ac in distinct ways across the three classes of POT peaks, revealing that Polycomb regulation of H3K27ac is not uniform across the genome. In PRC1cKO NGOs, H3K27ac levels were markedly elevated at “POT loss” peaks, whereas H3K27ac levels were reduced at “POT gain” peaks (Fig. 4C and fig. S4B). By contrast, in PRC2cKO NGOs, H3K27ac levels increased across all peak classes, consistent with the antagonistic relationship between H3K27me3 and H3K27ac (Fig. 4D and fig. S4B). In wild-type oocytes, H2AK119ub was gained at both “POT loss” peaks (Fig. 4E) and “POT gain” peaks (Fig. 4F) during POT, and PRC1 was required for both the loss and gain of H3K27ac at these sites. Together, these results indicate that PRC1 regulates H3K27ac in a site-selective manner.

In PRC1cKO NGOs, 17,460 H3K27ac peaks were differentially gained relative to PRC1ctrl NGOs, and these peaks were enriched for YY1 and MAFK motifs (Fig. 4, G and H). These motifs were also enriched among “POT loss” peaks in wild-type oocytes (Fig. 3C), supporting the idea that PRC1 facilitates developmental loss of H3K27ac during POT. Conversely, 4,749 H3K27ac peaks were differentially lost in PRC1cKO NGOs (Fig. 4G), and these peaks were enriched for TCF3 (E2A), PTF1A, and TCF12 (HEB) motifs (Fig. 4H), which were likewise enriched among “POT gain” peaks in wild-type oocytes (Fig. 3C). These findings support a second role for PRC1 in facilitating developmental gain of H3K27ac during POT. Thus, PRC1 influences both attenuation and acquisition of H3K27ac, indicating that it functions as more than a general suppressor of active chromatin.

Compared with PRC1 loss, PRC2 loss altered fewer peaks and produced a more uniform increase in H3K27ac signal (Fig. 4, D to G, and fig. S4B), consistent with the antagonism between H3K27me3 and H3K27ac at lysine 27. In PRC2cKO NGOs, both gained and lost H3K27ac peaks were similarly enriched for TCF3/12 motifs (Fig. 4H). Thus, PRC2 appears to contribute less selectively to H3K27ac redistribution during POT and primarily acts to restrain H3K27ac levels. The contrasting effects of PRC1cKO and PRC2cKO therefore reveal distinct Polycomb functions: PRC1 facilitates both developmental loss and gain of H3K27ac during POT, whereas PRC2-mediated H3K27me3 broadly antagonizes H3K27ac (Fig. 4I).

A prominent feature of FGOs is the presence of noncanonical, distal-rich broad H3K27ac domains (*18, 21*). To determine when and how these domains emerge, we examined H3K27ac enrichment across perinatal oocytes at genomic loci corresponding to all broad peaks (>10 kb) identified in FGOs (fig. S6A). Broad H3K27ac domains could already be detected in NGOs before growth, although they were relatively limited in early perinatal oocytes and became more evident during oocyte growth. Notably, in both PRC1- and PRC2-deficient NGOs, loci corresponding to broad H3K27ac domains in FGOs showed elevated H3K27ac relative to controls (fig. S6, A and B). These findings suggest that Polycomb normally restrains premature expansion of broad H3K27ac domains during ovarian reserve formation, further supporting a role for Polycomb in limiting untimely acquisition of active chromatin features in perinatal oocytes.

Together, these results indicate that PRC1 and PRC2 do not contribute equivalently to active chromatin remodeling during ovarian reserve formation. PRC2 broadly antagonizes H3K27ac, whereas PRC1 plays a more prominent and selective role in shaping developmental H3K27ac transitions during POT.

### Promoter bivalency is developmentally reconfigured during PPT

We next examined how promoter-associated chromatin states, particularly promoter bivalency, are regulated during perinatal oogenesis. Promoter bivalency, defined by the coexistence of H3K4me3 and H3K27me3, is typically associated with genes poised for later activation (*52*) and plays important roles in development and differentiation, particularly in the germline (*53, 54*). To define promoter bivalency, we classified promoters in E18O, NGOs, and GOs into four categories based on H3K4me3 and H3K27me3 status: bivalent, H3K27me3-only, H3K4me3-only, and other (fig. S7A). This analysis revealed that bivalent promoters were abundant at all three stages but underwent extensive developmental reconfiguration (Fig. 5A).

**Fig. 5.**
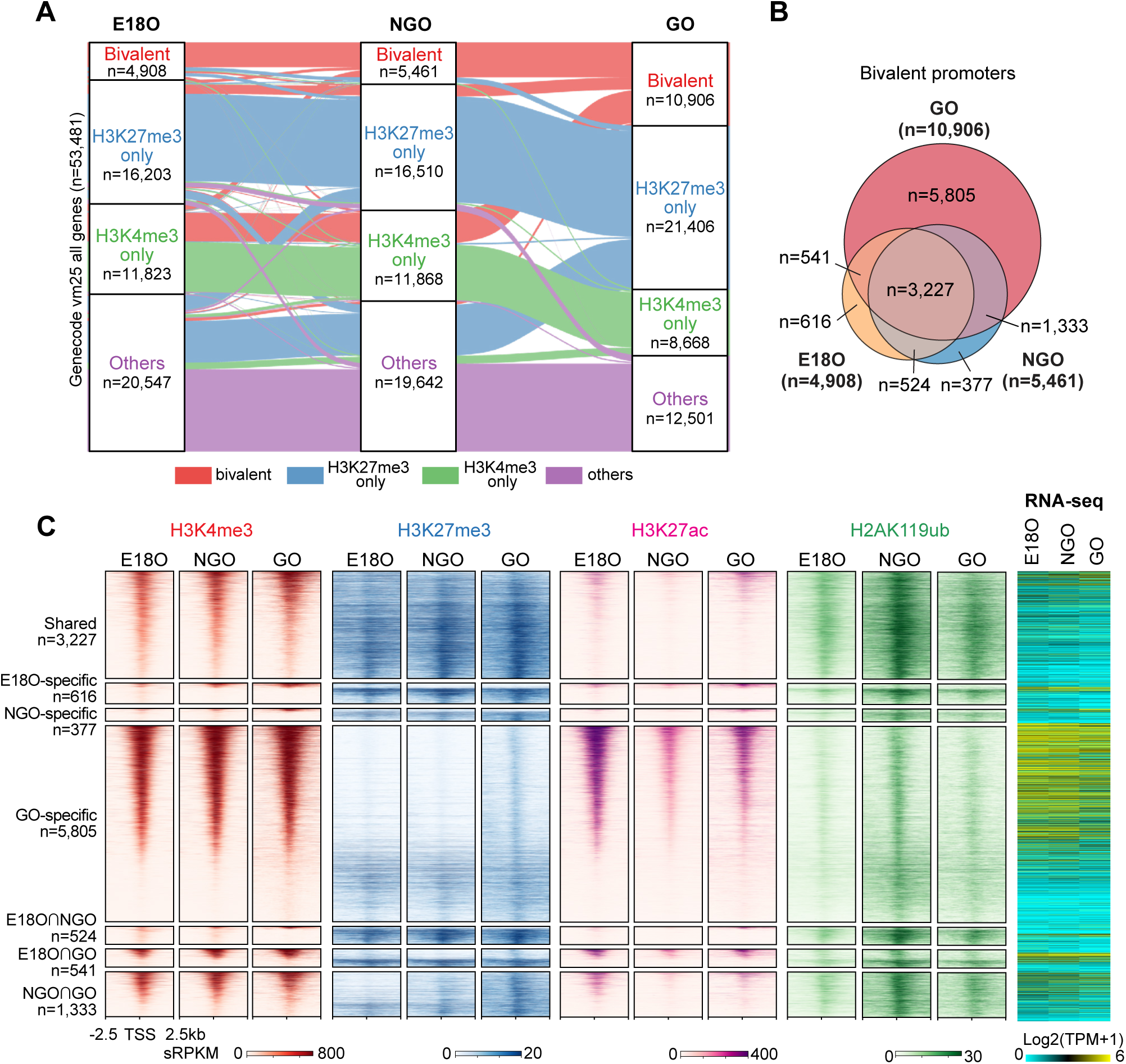
Dynamics of promoter bivalency during perinatal oogenesis. (A) Sankey diagram showing the distribution of bivalent, H3K27me3-only. H3K4me3-only, and other promoters in E18O, NGO and GO, respectively. Each promoter group in the GO stage is colored. (B) Venn diagrams showing bivalent promoter dynamics in perinatal oocytes. Numbers of total bivalent promoters in each stage are indicated. (C) Heatmaps showing changes in histone modifications at bivalent promoter regions (TSS ± 2.5 kb) in perinatal oocytes of each group indicated in (B). The color keys represent signal intensity, and the numbers represent sRPKM values. Right: Heat map showing RNA expression (log2-transformed TPM) of corresponding bivalent genes in WT oocytes.

A substantial fraction of bivalent promoters was shared across E18O, NGOs, and GOs, but many were newly established in GOs. We identified 4,908 and 5,461 bivalent promoters in E18Os and NGOs, respectively, with substantial overlap (3,751 in common), indicating that promoter bivalency is broadly maintained during POT (Fig. 5B and fig. S8A). By contrast, a substantial fraction of bivalent promoters detected in GOs (10,906 in total) arose from promoters that were not bivalent at earlier stages, particularly from H3K4me3-only promoters (Fig. 5B and fig. S8B), coincident with genome-wide H3K27me3 reprogramming during PPT (*30*). Because this reprogramming is directed by PRC1-H2AK119ub (*30*), H2AK119ub was already highly enriched at GO-specific bivalent promoters in NGOs (Fig. 5C), suggesting that H2AK119ub in NGOs primes the later formation of bivalent promoters in GOs.

Notably, virtually all classes of bivalent promoters were accompanied by H2AK119ub enrichment (Fig. 5C), indicating that these loci are more appropriately viewed as H2AK119ub-containing trivalent chromatin states and implicating PRC1 in regulation of promoter bivalency in oocytes. Among them, 3,227 promoters remained bivalent throughout all developmental stages and were marked by persistently high levels of H3K4me3 and H3K27me3 together with strong H2AK119ub (Fig. 5C). These constitutively bivalent promoters were associated with developmental regulators, canonical Polycomb targets, and morphogenesis-related processes (fig. S7B). By contrast, GO-specific bivalent promoters showed high H3K4me3 but relatively weak H3K27me3 and H2AK119ub enrichment, consistent with their higher expression levels than other classes of bivalent promoters (Fig. 5C and fig. S8B). Despite the relatively weak H3K27me3 enrichment, H3K27me3 deposition during PPT was still associated with reduced expression of these otherwise active genes (Fig. 5C and fig. S8B). Among these genes, those involved in translation, autophagy, and apoptosis were significantly down-regulated during PPT (fig. S7C).

Together, these findings indicate that promoter bivalency is dynamically reconfigured during perinatal oogenesis, with a major wave of de novo establishment during PPT. The presence of H2AK119ub at these promoters further suggests that many represent trivalent promoter states and implicates PRC1 in their regulation.

### Distinct roles of PRC1 and PRC2 at bivalent promoters during quiescence and early oocyte growth

Given the developmental reconfiguration of bivalent promoters during PPT, we next asked how Polycomb complexes regulate both the maintenance of preexisting bivalent promoters and the de novo establishment of new bivalent promoters during perinatal oogenesis. Because bivalent promoters are defined by the coexistence of the active mark H3K4me3 and the repressive mark H3K27me3, they provide a useful framework for dissecting how PRC1 and PRC2 intersect with promoter-associated active chromatin during POT and PPT.

To address this, we profiled H3K4me3 in PRC1cKO and PRC2cKO NGOs by quantitative CUT&Tag (fig. S9A). Unexpectedly, PRC1 and PRC2 deficiency had opposite effects on H3K4me3. PRC1 depletion significantly increased global H3K4me3 levels, particularly at peaks and promoter regions (fig. S9, B to G), whereas PRC2 deletion caused a modest genome-wide decrease in H3K4me3 (fig. S9, B to G). At bivalent promoters, PRC1 depletion increased H3K4me3 enrichment while leaving H3K27me3 largely unchanged, resulting in substantial derepression of associated genes in NGOs (Fig. 6A). Promoters that gained H3K4me3 in PRC1cKO NGOs largely overlapped with Polycomb-binding sites identified in other cell types (fig. S9H), consistent with H2AK119ub counteracting H3K4me3 at promoters (*29, 55*). By contrast, PRC2 deletion left the H2AK119ub profile at bivalent promoters unchanged, modestly reduced H3K4me3, and did not significantly alter expression of associated genes in NGOs (Fig. 6B). These results suggest that, in NGOs, H3K27me3 and H3K4me3 do not exhibit the antagonistic relationship observed in ESCs (*56*), and that PRC1 is the predominant regulator of bivalent promoters at the quiescent stage, consistent with the H2AK119ub-associated trivalent state observed in NGOs (Fig. 5C).

**Fig. 6.**
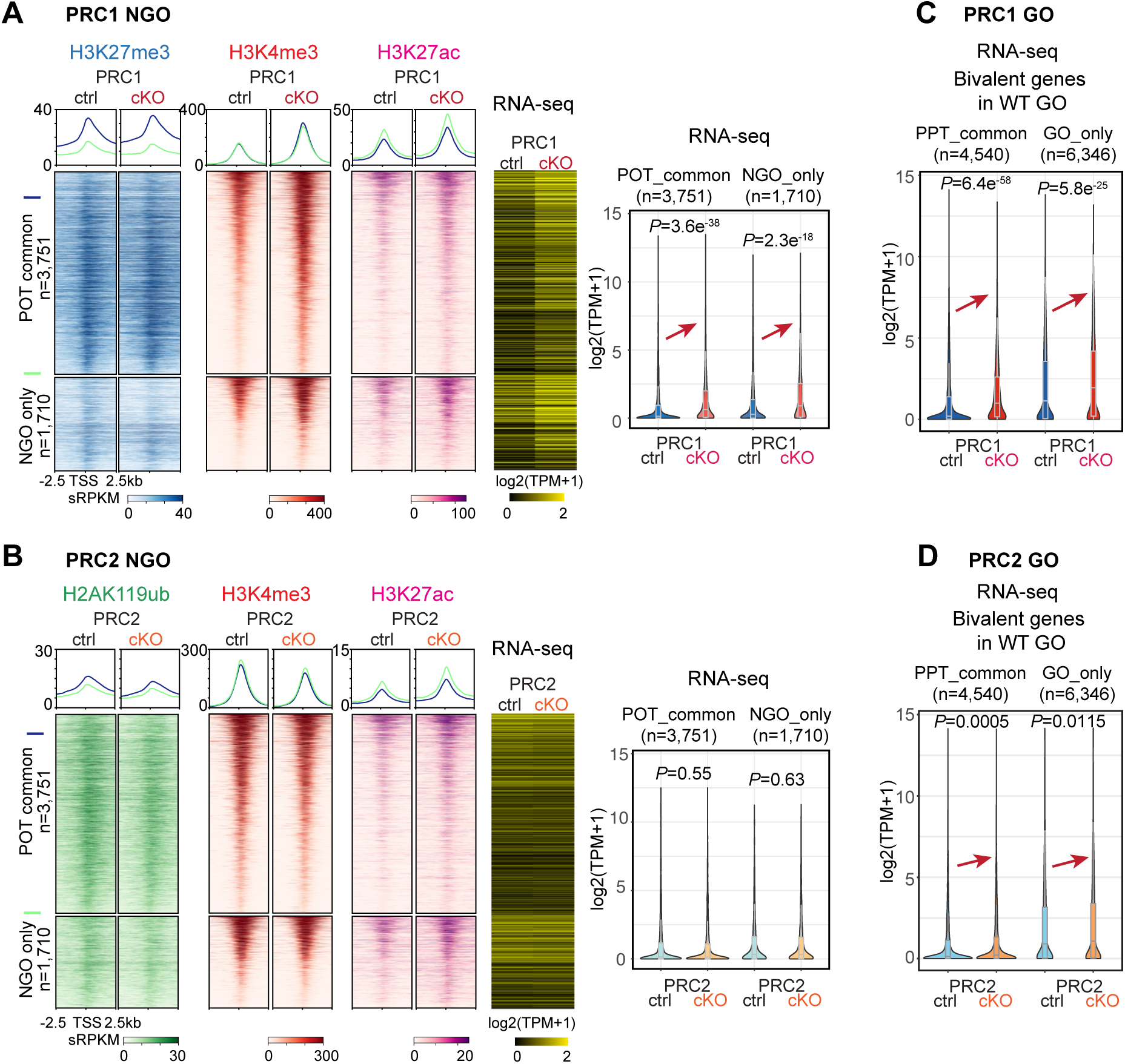
Distinct functions of PRC1 and PRC2 at bivalent promoters. (A and B) Average tag density plots and heatmaps showing histone modifications changes at WT NGO bivalent promoter regions (TSS ± 2.5 kb) in PRC1ctrl&cKO NGO (A) or PRC2ctrl&cKO NGO (B). The color keys represent signal intensity, and the numbers represent sRPKM values. Right: Heat map and violin plots with included boxplots showing RNA expression (log2-transformed TPM) of corresponding bivalent genes in PRC1ctrl and cKO NGO (A) or PRC2ctrl and cKO NGO (B). Boxes show the 25^th^ and 75^th^ percentile with the median, and whiskers indicate 1.5 times the interquartile range. *P* values of pairwise comparisons (two-sided unpaired Student’s t-test) are given. (C and D) Violin plots with included boxplots showing RNA expression (log2-transformed TPM) of corresponding bivalent genes in PRC1ctrl&cKO GO (C) or PRC2ctrl&cKO GO (D). Boxes show the 25^th^ and 75th percentile with the median, and whiskers indicate 1.5 times the interquartile range. *P* values of pairwise comparisons (two-sided unpaired Student’s t-test) are given.

This functional distinction between PRC1 and PRC2 was also evident in GOs. Genes associated with bivalent promoters in GOs, including both promoters maintained from NGOs and those newly established in GOs, were more strongly derepressed in PRC1-deficient GOs than in PRC2-deficient counterparts (Fig. 6, C and D). Notably, although loss of PRC2-H3K27me3 had little effect on expression of bivalent genes in NGOs (Fig. 6B), these genes tended to be up-regulated in GOs upon PRC2 loss, including genes shared between NGOs and GOs as well as newly established GO-specific bivalent genes (Fig. 6D). Thus, whereas PRC1 dominates restraint of bivalent promoters in NGOs, PRC2 begins to contribute more substantially to repression as oocytes enter growth.

Together, these findings reveal stage-dependent roles between PRC1 and PRC2 at bivalent promoters. PRC1 plays the stronger role in restraining H3K4me3 accumulation and transcription in NGOs, whereas PRC2 contributes more prominently after growth begins. More broadly, these results suggest that Polycomb-dependent regulation in perinatal oocytes helps couple long-term quiescence to future developmental competence by differentially controlling promoter-associated active chromatin states across the transition from the ovarian reserve to oocyte growth.

## Discussion

The ovarian reserve is the key determinant of female reproductive lifespan, yet the chromatin principles that stabilize NGOs while preserving competence for later growth have remained incompletely understood. Our previous work established that Polycomb-mediated repressive chromatin is integral to ovarian reserve formation and to later oocyte development (*14, 30*). Building on that framework, the present study uncovers an additional layer of Polycomb-dependent regulation centered on active chromatin (Fig. 7). H3K27ac is extensively redistributed during ovarian reserve formation, whereas both H3K27ac and H3K4me3 increase more broadly during subsequent oocyte growth. Moreover, PRC1 and PRC2 contribute unequally to these transitions, with PRC1 exerting the stronger influence on H3K27ac redistribution and promoter-associated H3K4me3 restraint. These findings indicate that the ovarian reserve is not an epigenetically inert state but is established through coordinated programming of both repressive and active chromatin.

**Fig. 7.**
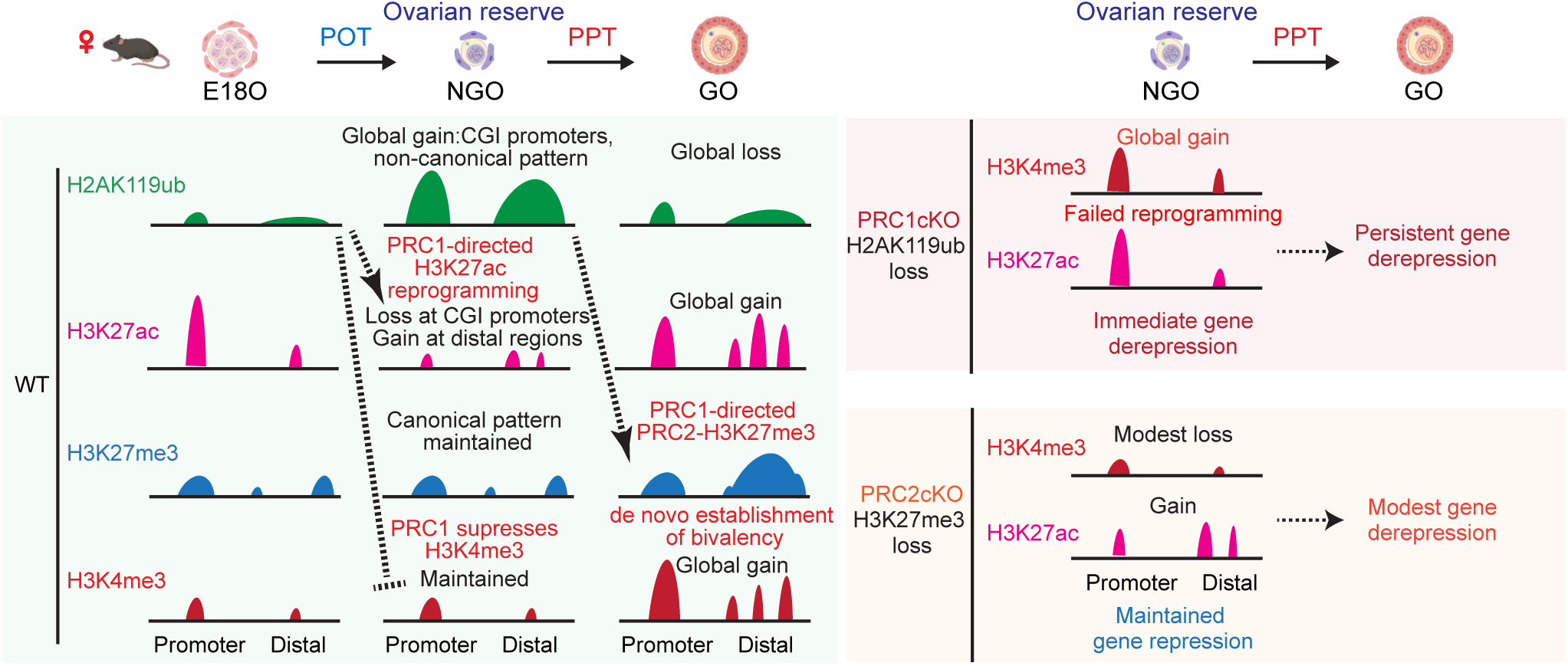
Model for Polycomb-directed crosstalk between active and repressive chromatin during perinatal oogenesis. Schematic model summarizing stage-specific crosstalk between Polycomb-mediated repressive chromatin and active chromatin during ovarian reserve formation and subsequent oocyte growth. During the perinatal oocyte transition, H3K27ac undergoes extensive redistribution, including CpG island promoter remodeling characterized by loss of H3K27ac and gain of PRC1-dependent H2AK119ub. This early promoter reprogramming is proposed to establish a repressive yet developmentally competent promoter framework in non-growing oocytes. During subsequent oocyte growth, broader gains in H3K27ac and H3K4me3 accompany acquisition of PRC2-dependent H3K27me3, de novo establishment of bivalent promoters, and formation of H2AK119ub-containing trivalent promoter states. PRC1 and PRC2 contribute unequally to these transitions: PRC2 broadly antagonizes H3K27ac, whereas PRC1 plays the stronger role in shaping developmental H3K27ac remodeling and restraining promoter-associated H3K4me3. Together, these processes establish promoter states that support quiescence while preserving future growth potential and developmental competence.

A central implication of our data is that CGI promoter reprogramming during ovarian reserve formation may help preconfigure promoter states for later oocyte growth and embryogenesis. During this period, many CGI promoters lose H3K27ac while gaining PRC1-mediated H2AK119ub, regardless of whether the associated genes are subsequently up- or down-regulated. This promoter-centered remodeling is followed during oocyte growth by acquisition of PRC2-mediated H3K27me3 and de novo establishment of bivalent promoters, supporting a model in which early promoter reprogramming creates a chromatin framework for later promoter-state transitions. This model is especially relevant in the context of DNA methylation establishment during oocyte growth. In growing oocytes, de novo DNA methylation is deposited broadly across transcribed regions, whereas CGI promoters generally remain protected from methylation (*38*). Loss of this protection causes persistent maternal CGI hypermethylation, impaired embryonic gene expression, and preimplantation lethality (*57*). In this context, our findings raise the possibility that early CGI promoter remodeling in NGOs, characterized by loss of H3K27ac and gain of Polycomb-associated marks, helps establish a promoter chromatin state that is less permissive to inappropriate de novo DNA methylation during later oocyte growth, thereby contributing to formation of the maternal epigenome inherited by the next generation. In this sense, perinatal oogenesis is better viewed not as simple entry into quiescence, but as a developmental window in which promoter states are reorganized in preparation for future oocyte development.

Our data further suggest that these later poised promoter states extend beyond the classical PRC2-centered concept of bivalency. Rather than representing classical bivalent domains marked only by H3K4me3 and H3K27me3, a substantial fraction of these poised promoters in oocytes may be more appropriately viewed as trivalent promoter states because they are also enriched for H2AK119ub. This is notable because it places PRC1-H2AK119ub directly within the chromatin architecture of poised developmental promoters in oocytes. Consistent with this interpretation, PRC1 loss had a stronger effect than PRC2 loss on promoter-associated H3K4me3 accumulation, indicating that PRC1-H2AK119ub is an important determinant of H3K4me3 restraint during establishment and maintenance of these bivalent/trivalent promoter states. Thus, PRC1 appears to act not only upstream of later H3K27me3 accumulation, but also as a key regulator of promoter chromatin balance during ovarian reserve formation and subsequent growth.

Our perturbation analyses further show that PRC1 and PRC2 do not act equivalently in the active chromatin remodeling process. PRC2 broadly antagonizes H3K27ac, consistent with the mutually exclusive nature of H3K27ac and H3K27me3 (*58–63*). PRC1, by contrast, acts more selectively, facilitating both developmental loss and gain of H3K27ac at distinct loci. Previous studies have shown that PRC1 can counteract H3K27ac by inhibiting acetyltransferase recruitment (*44*) or recruiting deacetylases (*45*), but in some contexts it can also support gene activation and H3K27ac accumulation (*64–67*). Our data are consistent with such context-dependent regulation and suggest that PRC1 functions not simply as a general suppressor of active chromatin, but as a selective organizer of promoter remodeling during ovarian reserve formation. We further find that broad H3K27ac domains characteristic of FGOs can already be detected in NGOs and become more prominent during growth, while both PRC1 and PRC2 restrain their premature expansion during ovarian reserve formation. Together, these observations support a broader role for Polycomb in delimiting the timing and extent of active chromatin acquisition.

This interpretation also helps unify our recent studies. Our previous work showed that epigenetic priming in NGOs shapes later chromatin states during oocyte growth and the oocyte-to-embryo transition, and that PRC1-dependent H2AK119ub prepatterns later H3K27me3 organization and developmental competence (14, 30). The present study adds the missing promoter-level logic to this framework: PRC1-dependent CGI promoter remodeling during the perinatal stage, especially removal of H3K27ac coupled to gain of H2AK119ub, appears to create chromatin states that are later elaborated into bivalent/trivalent promoters and protected from inappropriate DNA methylation during growth. In this way, the current study supports a broader model in which epigenetic priming in the female germline is established through staged Polycomb-dependent remodeling of both active and repressive chromatin.

More broadly, our findings place perinatal oocytes within an emerging view of Polycomb regulation in which poised promoter states are not simply inherited, but developmentally assembled and reconfigured. In pluripotent cells, bivalent promoters have classically been interpreted through a PRC2-centered framework, in which H3K27me3 counterbalances promoter-associated H3K4me3 to maintain developmental genes in a poised state (*52–54*). Our data suggest that perinatal oocytes deploy a germline-adapted version of this logic, in which PRC1 plays a disproportionately strong role at an earlier stage by remodeling CGI promoters during ovarian reserve formation and limiting H3K4me3 accumulation and transcriptional leakage at later poised promoters. At the same time, although the H3K27ac and H3K4me3 changes described here are tightly associated with developmental progression and gene expression changes, our study does not support a generalized conclusion that such chromatin changes directly determine all aspects of later developmental competence. Resolving causality will require targeted functional dissection of individual loci and regulatory elements.

In summary, our study identifies perinatal oogenesis as a developmental window in which Polycomb-mediated repressive programming and active chromatin remodeling are tightly integrated (Fig. 7). During this period, PRC1-dependent CGI promoter reprogramming is coupled to H3K27ac redistribution and bivalent promoter priming, thereby helping establish promoter states that support quiescence while preserving future growth potential. These findings extend the current framework of ovarian reserve biology beyond Polycomb-dependent repression, establish active chromatin remodeling as an integral component of perinatal oocyte development, and provide a molecular logic for how epigenetic priming is established in the female germline.

## Methods

### Animals

The oocyte-specific reporter *Stella*-GFP transgenic mice were obtained from Dr. M. Azim Surani (*68*). Generation of conditionally deficient *Rnf2* mice on a *Ring1^-/-^*background with *Ddx4-Cre* was performed as described previously (*14, 46, 47*). Briefly, PRC1*cKO* mice *Ring1^-/-^; Ring1^F/-^; Ddx4-Cre* were generated from *Ring1^-/-^; Ring1^F/F^* females crossed with *Ring1^-/-^; Ring1^F/+^; Ddx4-Cre* males and PRC1*ctrl* mice used in experiments were *Ring1^-/-^; Rnf2^F/+^; Ddx4-Cre* littermate females. PRC2*cKO* mice *Eed^F/-^; Ddx4-Cre* were generated from *Eed^F/F^* females crossed with *Eed^F/+^; Ddx4-Cre* males and *PRC2ctrl* mice used in experiments were *Eed^F/+^; Ddx4-Cre* littermate females. *Eed^F/F^* mice were also crossed with the *Stella*-GFP line to get both PRC2*ctrl* and *cKO* mice with the GFP reporter for oocyte collection using FACS. Generation of mutant *Ring1* and *Rnf2* floxed alleles were reported previously (*50, 69*). The *Eed flox* (B6;129S1-*Eed* ^tm1Sho^/J) and *Ddx4-Cre* [FVB-Tg(Ddx4-cre)1Dcas/J] mouse lines were purchased from the Jackson Laboratory (022727, 006954 and 011062, respectively). Mice were maintained on a mixed genetic background of FVB and C57BL/6J.

Mice were maintained on a 12:12 light: dark cycle in a temperature and humidity-controlled vivarium (22 ± 2 °C; 40–50% humidity) with free access to food and water in the pathogen-free animal care facility. Mice were used according to the guidelines of the Institutional Animal Care and Use Committee (IACUC protocol no. 21931) at the University of California, Davis.

### Oocyte collection

For WT profiling experiments, small oocytes (E18.5, P1, P6) from *Stella*-GFP reporter female mice were isolated by FACS and the early GOs (P7) were manually collected under the stereomicroscope. For PRC1 conditional knockout studies, both NGOs and GOs were manually collected using a stereomicroscope. For PRC2 conditional knockout studies, NGOs were isolated from PRC2ctrl or cKO female pups carrying *Stella*-GFP reporter by FACS and GOs were manually collected using a stereomicroscope.

Ovaries at different developmental stages were harvested by carefully removing oviducts and ovarian bursa in PBS. Each pair of ovaries were further digested in 200 μl TrypLE™ Express Enzyme (1X) (Gibco, 12604013) supplemented with 0.3% Collagenase Type 1 (Worthington, CLS-1) and 0.01% DNase I (Sigma, D5025) and incubated at 37℃ for 20-40 min (longer time for bigger postnatal ovaries) with gentle agitation. After incubation, the ovaries were dissociated by gentle pipetting using the Fisherbrand^TM^ Premium Plus MultiFlex Gel-Loading Tips until no visible tissue pieces. Two ml DMEM/F-12 medium (Gibco, 11330107) supplemented with 10% FBS (HyClone, SH30396.03) were then added to the suspension to stop enzyme reaction.

For FACS preparation, the cells were suspended in FACS buffer (PBS containing 2%FBS) and subjected to centrifugation at 300 × g for 5 min. The supernatant was decanted and the cells resuspended in FACS buffer and filtered into a 5 ml FACS tube with a 35 μm nylon mesh cap (Falcon, 352235). The cells were analyzed after removing small and large debris in FSC-A versus SSC-A gating, and doublets in FSC-W versus FSC-H gating. Then, the desired small oocyte population (enriched GFP+ cells with uniform size) was collected based on both GFP signal gating and FSC-A versus SSC-A size backgating.

For manual picking, the cell suspension was filtered through a 100 μm cell strainer and seeded onto a culture dish. The cells were allowed to settle down for 15-30 min at 37℃ in an atmosphere containing 5% CO_2_ in air before being transferred under a stereomicroscope (Nikon, SMZ1270). Based on morphology and diameter, non-growing oocytes or growing oocytes were specifically picked up, washed, and transferred into the downstream buffer with a mouth-operated glass capillary pipette.

### Quantitative CUT&Tag library generation and sequencing

CUT&Tag libraries of oocytes were prepared as previously described (*31, 32*) with some modifications (a step-by-step protocol https://www.protocols.io/view/bench-top-cut-amp-tag-kqdg34qdpl25/v3) using CUTANA™ pAG-Tn5 (Epicypher, 15-1017). To perform quantitative spike-in CUT&Tag, Drosophila S2 cells were added to mouse oocytes at a fixed ratio (e.g., 2000 S2 cells to 1000 mouse oocytes) at the beginning of each reaction. The antibodies used were rabbit anti-H3K4me3 (1/50; Active Motif; 39159) and rabbit anti-H3K27ac (1/50; Cell Signaling Technology; 8173). CUT&Tag libraries were sequenced on the HiSeq 4000 system (Illumina) with 150-bp paired-end reads.

### CUT&Tag data processing

#### Upstream processing

We basically followed the online tutorial posted by the Henikoff Lab (https://protocols.io/view/cut-amp-tag-data-processing-and-analysis-tutorial-bjk2kkye.html) with some modifications. Briefly, after trimming by Trim-galore (https://github.com/FelixKrueger/TrimGalore) (version 0.6.10), raw paired-end reads were aligned with the mouse genome (GRCm38/mm10) using Bowtie2 (*70*) (version 2.4.5) with options: --end-to-end --very-sensitive --no-mixed --no-discordant --phred33 -I 10 -X 700. *D. melanogaster* DNA delivered by *Drosophila* S2 cells was used as spike-in DNA for CUT&Tag. For mapping *D. melanogaster* spike-in fragments, we also use the --no-overlap --no-dovetail options to avoid cross-mapping. PCR duplicates from all mapped bam files were removed using the ‘MarkDuplicates’ command in Picard tools (version 3.0.0) (https://broadinstitute.github.io/picard/). To compare replicates, Pearson correlation coefficients were calculated and plotted by ‘multiBamSummary bins’ and ‘plot correlation’ commands from deepTools (*71*) (version 3.5.5). Biological replicates were pooled using the ‘merge’ command in samtools for visualization and other analyses after confirming reproducibility (Pearson correlation coefficient > 0.85). Spike-in normalization was implemented using the exogenous scaling factor computed from the dm6 mapping files (scale factors = 10,000 / aligned spike-in reads).

#### Downstream analysis and visualization

Spike-in normalized genome coverage tracks with 1 bp resolution in BigWig format were generated using ‘bamCoverage’ from deepTools with the parameters ‘--binSize 1 --extendReads --samFlagInclude 64 --normalizeUsing RPKM --scaleFactor $scale_factor’. Track views were visualized and exported from Integrative Genomics Viewer (*72*) (version 2.14.1). Average scores (sRPKM: scaled RPKM) per region [10kb bins or custom intervals, e.g., peaks, promoters (TSS±1kb)] were calculated from scaled BigWig files using the ‘multiBigwigSummary bins’ or ‘multiBigwigSummary BED-file’ functions from deepTools with adding parameter ‘--outRawCounts’. Blacklisted regions were excluded from analyses by applying the parameter ‘--blackListFileName mm10-blacklist.v2.bed’ while using deepTools. This output matrix from deepTools is used for the following comparative analyses. Hierarchical clustering analysis was conducted in Morpheus using sRPKM values of each 10kb bin across the entire genome as input. Sum sRPKM values of all bins from each replicate track were used to calculate relative global levels of histone modifications. Following a previous study (*73*), average sRPKM values of 10kb bins or custom intervals are transformed to integers and used as input for differential enrichment analyses, performed by DESeq2 with fixed size factors since BigWig files were already spike-in normalized. Differential enriched regions are identified with cutoffs log2FoldChange > 1 and P_adj_ < 0.05. Plots were created using the R package ggplot2 (*74*) (v3.5.1).

Bivalent promoters were characterized based on the enrichment level (sRPKM values) of H3K4me3 and H3K27me3 on promoter regions (TSS±1kb) calculated by deepTools. All the promoters (H3K4me3 sRPKM>20 & H3K27me3 sRPKM>2) were defined as bivalent promoters. The cutoff values were confirmed by manual visualization of the track views.

MACS2 (*75*) was used for peak calling of H3K4me3 and H3K27ac with the parameters’-p 1e-5 - -broad -g mm’. The H3K27ac peaks were categorized into proximal peaks (TSS ±1 kb) and distal peaks (beyond TSS ±1 kb). HOMER (*76*) (version 4.11) was used for peak annotation and finding motifs. Peaks comparison was done by the ‘intersect’ command of bedtools (*77*) (version 2.31.0). For differential enrichment analyses on peaks, peaks of each genotype or stage were merged by BEDtools and then used as the reference genomic intervals for counting. ChIP-x Enrichment Analysis (ChEA) was performed using the Enricher website (https://maayanlab.cloud/Enrichr/) (*78, 79*). DeepTools was used to draw tag density plots and heatmaps for enrichments.

### Chromatin states analysis using ChromHMM

The ChromHMM (*33*) software (version 1.25) was used to classify genomic regions based on the dynamics of all four histone modifications in perinatal oogenesis. Specifically, signal files in bed format (transformed from above scaled Bigwig files) for each histone modification from all stages were first binarized using the ‘BinarizeBed’ command with a bin size of 5 kb (option ‘-b 5000’). The binarized signal files were then used as input to characterize chromatin states conducted using the ‘LearnModel’ command with 5-kb bins (options’-b 5000’). The number of states to include in the mode was finally set to 12 for best visualization after comparing outputs for different numbers.

### Identification of CGI genes and non-CGI genes

CpG islands (CGIs) were defined as published (*80, 81*), and their coordinates of mouse genome GRCm38/mm10 assembly (n=17,017) were downloaded from UCSC genome browser (https://genome.ucsc.edu/cgi-bin/hgTables?db=mm10&hgta_group=regulation&hgta_track=cpgIslandExt&hgta_table=cpgIslandExt&hgta_doSchema=describe+table+schema). Genes were classified as CGI-associated if a first base pair of any transcript TSSs ±100-bp overlapped a CGI. In this way, the total 53,795 genes from Gencode vM25 mouse gene annotations (after removing Chromosome Y and M genes) were divided into CGI genes (n=14,040) and non-CGI genes (n=39,755).

### RNA-seq data reanalysis

Raw RNA-seq datasets of WT oocytes were downloaded from GSE128305 (ref.(*11*)). Raw RNA-seq reads after trimming by Trim-galore (https://github.com/FelixKrueger/TrimGalore) (version 0.6.10) were aligned to the mouse (GRCm38/mm10) genome using HISAT2 (*82*) (version 2.2.1) with default parameters. All unmapped reads, non-uniquely mapped reads, and reads with low mapping quality (MAPQ < 30) were filtered out by samtools (*83*) (version 1.16.1) with the parameter ‘-q 30’ before being subjected to downstream analyses. To identify differentially expressed genes in RNA-seq, raw read counts for each gene were generated using the featureCounts(*84*) part of the Subread package (version 2.0.3) based on mouse gene annotations (gencode.vM25.annotation.gtf, GRCm38/mm10). DESeq2 (*85*) (version 1.44.0) was used for differential gene expression analyses with cutoffs FoldChange > 2 and P_adj_ values < 0.05.

GO analyses were performed using the online functional annotation clustering tool Metascape (*86*) (http://metascape.org). The TPM values of each gene were generated using RSEM (*87*)(version 1.3.3) for comparative expression analyses and computing the Pearson correlation coefficient between biological replicates. RNA expression heatmaps were plotted using Morpheus (https://software.broadinstitute.org/morpheus/).

### ChIP-seq data reanalysis

Raw STAR ChIP-seq data of H3K27ac in GV FGOs were downloaded from Gene Expression Omnibus (GEO) under accession no. GSE217970 (ref.(*18*)). Raw reads files were processed using the same pipeline as CUT&Tag datasets for alignment and duplicate removal. Deduped bam files were converted to bigwig files using deeptools by normalizing with RPKM and then used for plotting. Peak calling for H3K27ac was performed using the SEACR (*88*) (https://seacr.fredhutch.org/) by selecting the top 5% enriched regions in stringent mode with parameters’-c 0.05 -n norm -m stringent’. Peaks (width>10kb) were selected as broad domains and used for plotting.

### WGBS data reanalysis

Raw WGBS data of GV FGOs were downloaded from DDBJ: DRA000570 (ref. (*15*)). Low-quality bases at the 3’ end and residual adapter sequences were removed from the raw sequencing reads. The processed reads were then aligned to the mouse genome (mm10) using Bismark with the PBAT option. Bisulfite conversion rates were estimated from reads uniquely aligned to the lambda phage genome. Counts from the two cytosines in a CpG dinucleotide and its reverse complement were combined, and only CpG sites with coverage between 5× and 200× were included in the analysis of CpG methylation levels.

### Statistics

Statistical methods and P values for each plot are listed in the figure legends and/or in the Methods. Statistical significance for summary quantifications was determined using two-tailed unpaired t-tests. Next-generation sequencing data (RNA-seq, CUT&Tag) were based on two independent replicates. No statistical methods were used to initially determine sample size in these experiments. Experiments were not randomized, and investigators were not blinded to allocation during experiments and outcome assessments.

### Data availability

RNA-seq datasets from PRC1 ctrl&cKO and PRC2 ctrl&cKO oocytes were first reported in (*14, 30*) under accession nos. GSE280638 and GSE184208, respectively. RNA-seq datasets of WT oocytes were downloaded from GSE128305 (*11*). ATAC-seq data were obtained from GSE306919 (*89*). Raw STAR ChIP-seq data of H3K27ac in GV FGOs were downloaded from GSE217970 (*18*). The DNA methylation WGBS data of GV full-grown oocytes were from DDBJ: DRA000570 (*15*).

### Code availability

Source code for all software and tools used in this study, with documentation, examples, and additional information, is available at the URLs listed above. No custom code was generated beyond standard public tools.

## Acknowledgments

We thank M. Azim Surani for sharing *Stella*-GFP transgenic mice, and Yao Cai and Joanna Chiu for sharing Drosophila S2 cells. Funding sources: Open Collective Foundation Repro Grants (M.H), National Institutes of Health grants R35GM141085 (S.H.N) and R21HD110146 (S.H.N and R.M.S).

## Author contributions

M.H. and S.H.N. designed the study. M.H., Y.H.Y. performed experiments. M.H., Y.M. and S.H.N. designed and interpreted the computational analyses. M.H., N.H., R.M.S., and S.H.N. interpreted the results and wrote the manuscript with critical feedback from all other authors. R.M.S. and S.H.N. supervised the project.

