## Supplementary Figures for "Polycomb shapes active chromatin and promoter bivalency during ovarian reserve formation and activation"

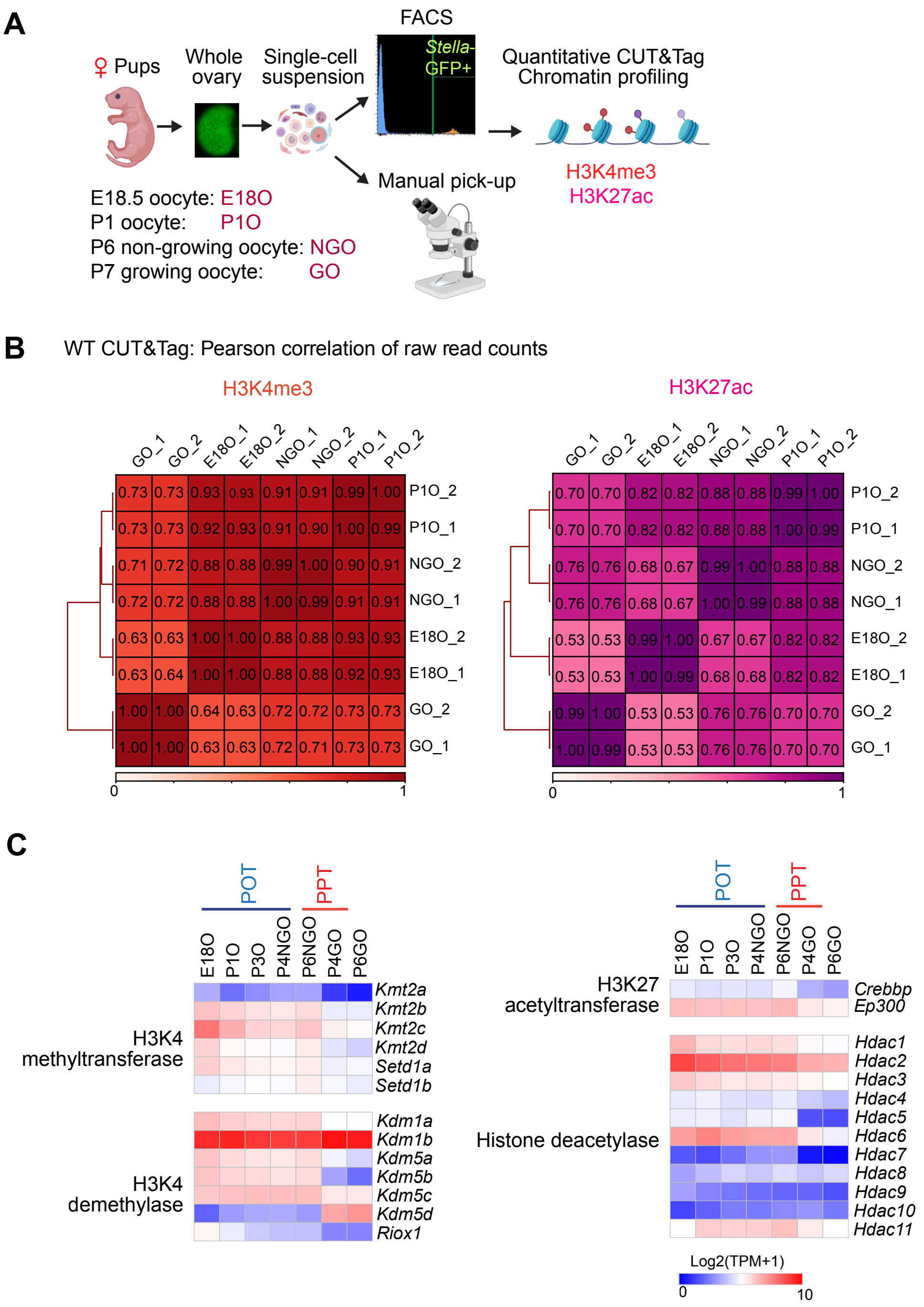


**Fig. S1. Quantitative CUT&Tag workflow, data quality, and expression of active histone mark regulators during perinatal oogenesis.**

(A) Overview of the experimental steps for chromatin profiling using quantitative CUT& Tag on perinatal oocytes (created with BioRender.com).

(B) Heatmaps with hierarchical clustering showing the Pearson correlation of the raw read counts among each biological replicate in WT CUT&Tag data.

(C) Heatmap showing gene expression of the key components comprising enzymes related to H3K4me3 and H3K27ac during perinatal oogenesis. In wild-type, E18O represents oocytes in MPI; P1O and P3O represent oocytes transitioning to dictyate arrest;P4 and P6 small oocytes represent NGOs residing in primordial follicles; and P4 and P6 large oocytes represent GOs in primary follicles after initiation of oocyte growth. RNA-seq data of WT oocytes were downloaded from GSE128305 (ref.[11]) and processed. TPM values of genes were used to plot the heatmaps.

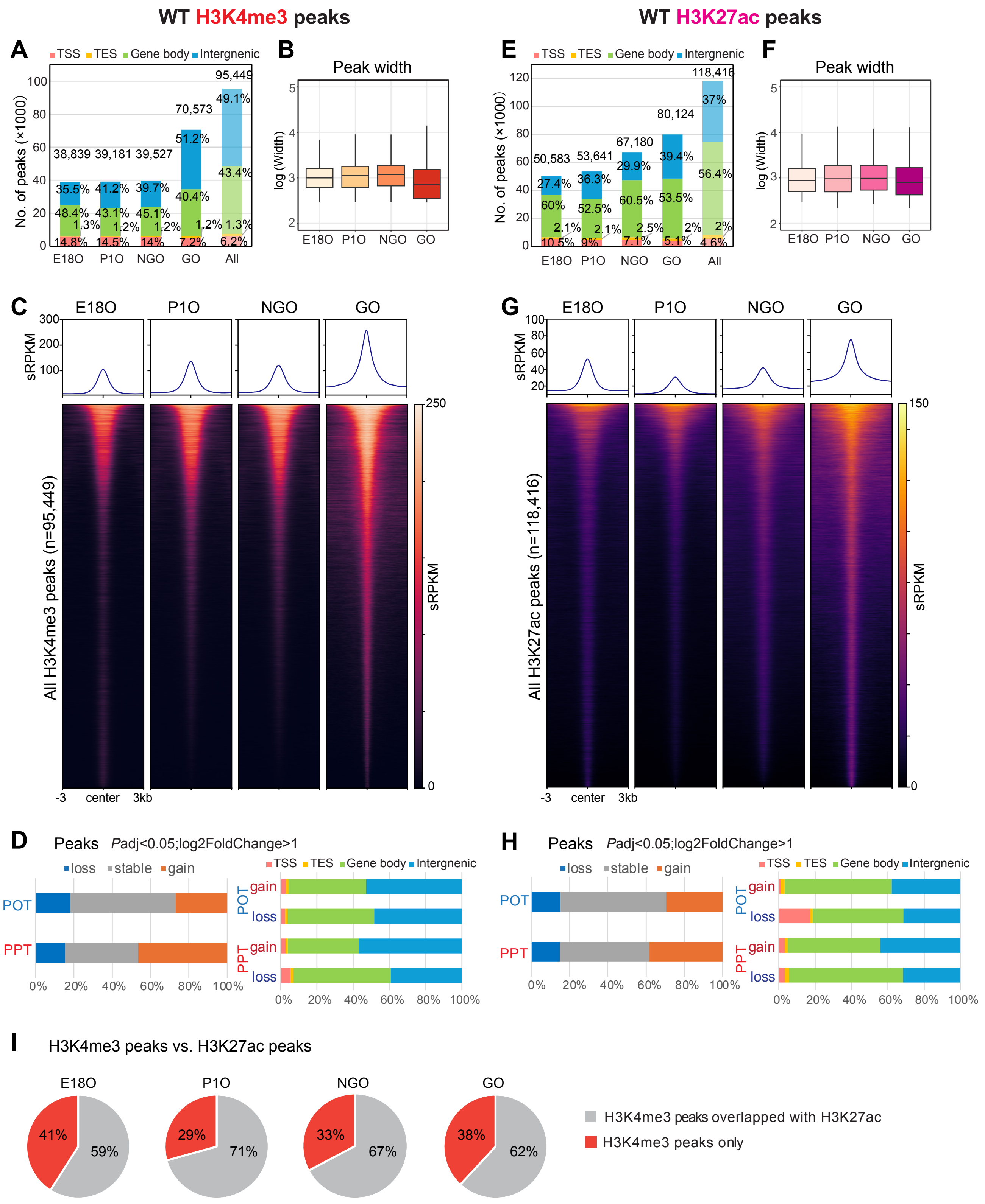
WT H3K4me3 peaks
WT H3K27ac peaks
A
TSS TES Gene body Intergenic
No. of peaks (×1000)
38,839 39,181 39,527 70,573 95,449
35.5% 41.2% 39.7% 51.2% 49.1%
48.4% 43.1% 45.1% 40.4% 43.4%
1.3% 1.2% 1.2% 1.2% 1.3%
14.8% 14.5% 14% 7.2% 6.2%
E18O P1O NGO GO All
B
Peak width
log (Width)
E18O P1O NGO GO
E
TSS TES Gene body Intergenic
No. of peaks (×1000)
50,583 53,641 67,180 80,124 118,416
27.4% 36.3% 29.9% 39.4% 37%
60% 52.5% 60.5% 53.5% 56.4%
2.1% 2.1% 2.5% 2% 2%
10.5% 9% 7.1% 5.1% 4.6%
E18O P1O NGO GO All
F
Peak width
log (Width)
E18O P1O NGO GO
C
E18O P1O NGO GO
sRPKM
All H3K4me3 peaks (n=95,449)
-3 center 3kb
G
E18O P1O NGO GO
sRPKM
All H3K27ac peaks (n=118,416)
-3 center 3kb
D
Peaks Padj<0.05;log2FoldChange>1
loss stable gain
POT
PPT
TSS TES Gene body Intergenic
gain
loss
0% 20% 40% 60% 80% 100%
H
Peaks Padj<0.05;log2FoldChange>1
loss stable gain
POT
PPT
TSS TES Gene body Intergenic
gain
loss
0% 20% 40% 60% 80% 100%
I
H3K4me3 peaks vs. H3K27ac peaks
E18O P1O NGO GO
41% 59%
29% 71%
33% 67%
38% 62%
H3K4me3 peaks overlapped with H3K27ac
H3K4me3 peaks only

**Fig. S2. Dynamics of active marks H3K4me3 and H3K27ac during perinatal oogenesis.**
(A, E) Bar charts showing H3K4me3 and H3K27ac peak number and genomic distribution in WT perinatal oocytes.
(B, F) Boxplots showing H3K4me3 and H3K27ac peak width in WT perinatal oocytes. Boxes show the 25th and 75th percentile with the median, and whiskers indicate 1.5 times the interquartile range.
(C, G) Heatmaps and average tag density plots showing all H3K4me3 and H3K27ac peaks in WT perinatal oocytes.
(D, H) Bar charts showing percentages of differentially enriched peaks for H3K4me3 and H3K27ac during POT and PPT and their corresponding genomic distribution.
(I) Pie chart showing percentages of H3K4me3 peaks overlapping H3K27ac peaks in each stage of perinatal oogenesis.

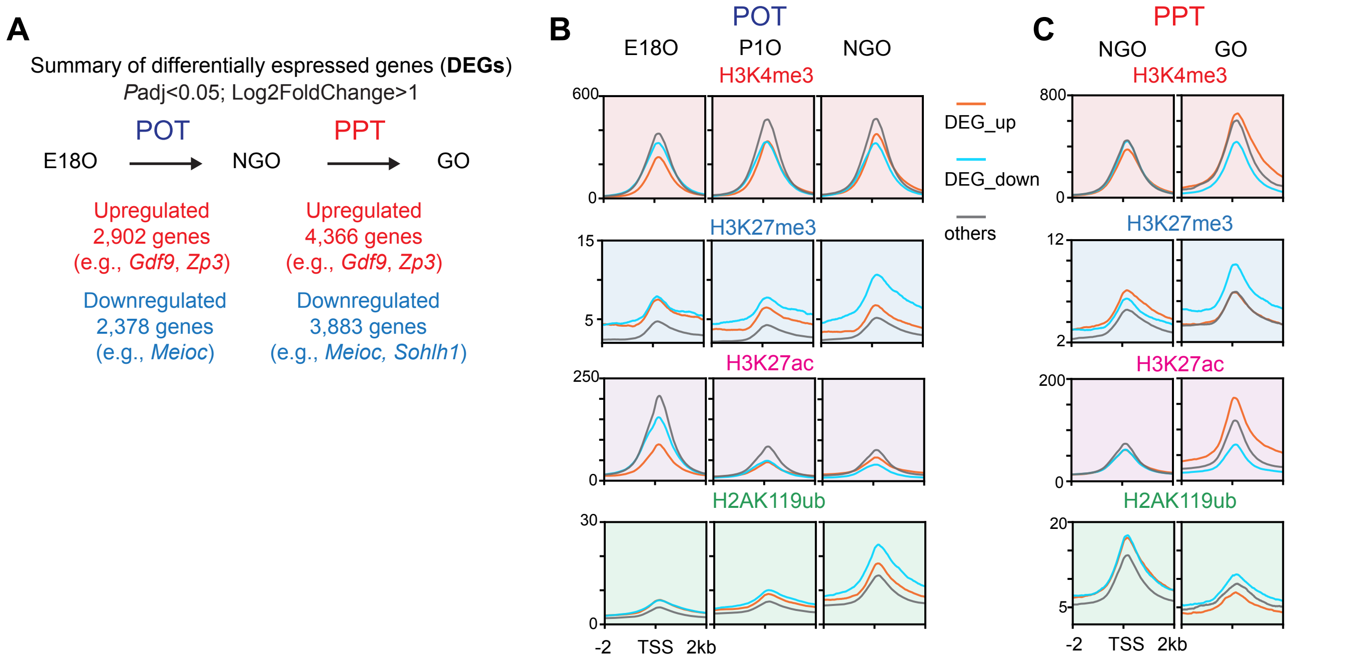


**Fig. S3. Histone modifications at promoters of differentially expressed genes during POT and PPT.** (A) Summary of differentially expressed genes (DEGs) during POT and PPT identified in WT oocytes. (B, C) Average tag density plots showing H3K4me3, H3K27me3, H3K27ac and H2AK119ub enrichment at promoter regions (TSS ± 2 kb) of DEGs during POT and PPT in WT mice.

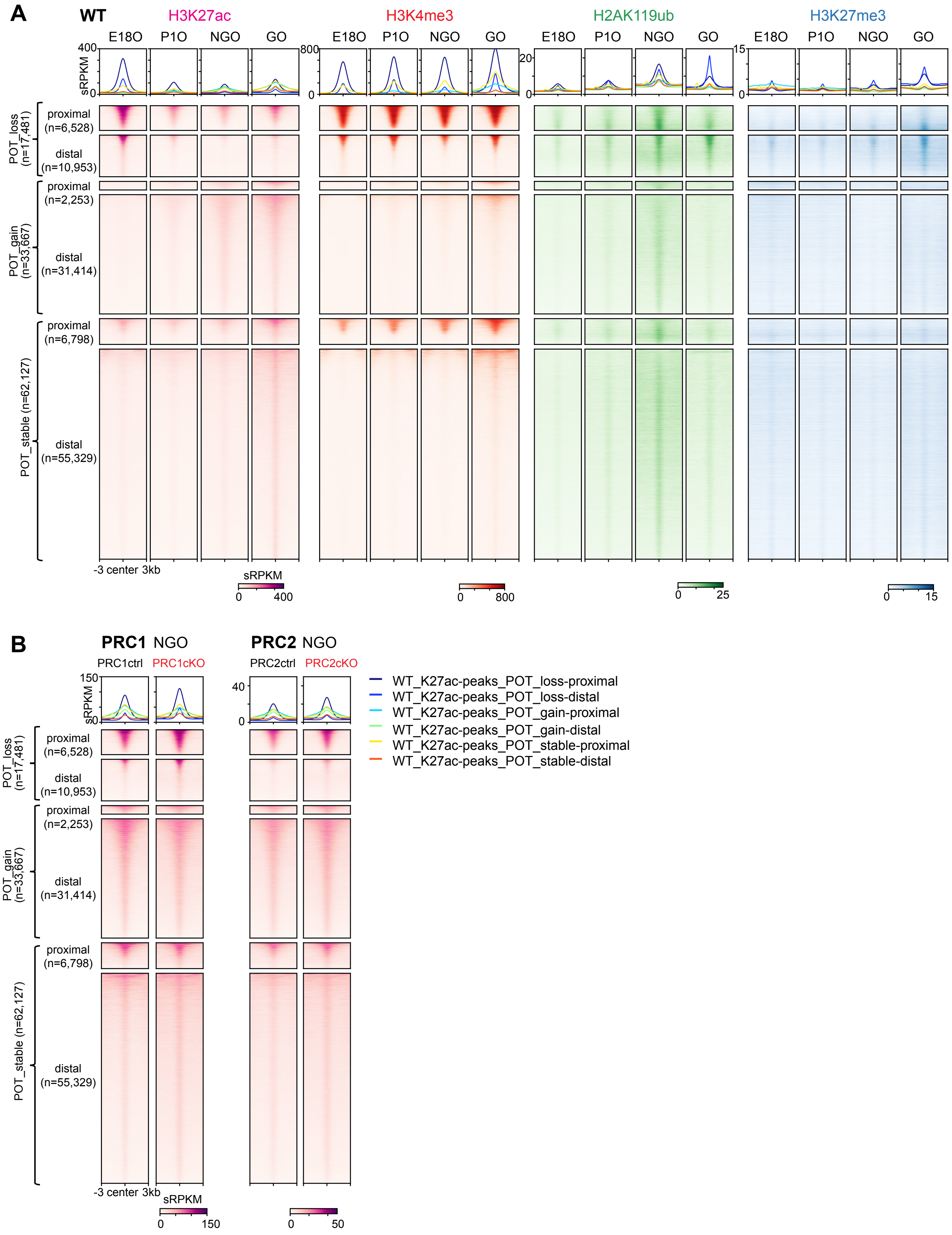
A
WT
H3K27ac
H3K4me3
H2AK119ub
H3K27me3
E18O
P1O
NGO
GO
sRPKM
400
800
20
15
0
POT_loss (n=17,481)
proximal (n=6,528)
distal (n=10,953)
POT_gain (n=33,667)
proximal (n=2,253)
distal (n=31,414)
POT_stable (n=62,127)
proximal (n=6,798)
distal (n=55,329)
-3 center 3kb
sRPKM
0
400
0
800
0
25
0
15
B
PRC1 NGO
PRC2 NGO
PRC1ctrl
PRC1cKO
PRC2ctrl
PRC2cKO
150
40
sRPKM
WT_K27ac-peaks_POT_loss-proximal
WT_K27ac-peaks_POT_loss-distal
WT_K27ac-peaks_POT_gain-proximal
WT_K27ac-peaks_POT_gain-distal
WT_K27ac-peaks_POT_stable-proximal
WT_K27ac-peaks_POT_stable-distal
-3 center 3kb
sRPKM
0
150
0
50

**Fig. S4. Histone modification at proximal and distal regions of differential H3K27ac peaks.** (A, B) Average tag density plots and heatmaps showing histone modification changes at proximal and distal regions of differential H3K27ac peaks (peak center ± 3 kb) in E18O and NGO of WT (A), and NGO of PRC1ctr&cKO and PRC2ctrl&cKO (B).

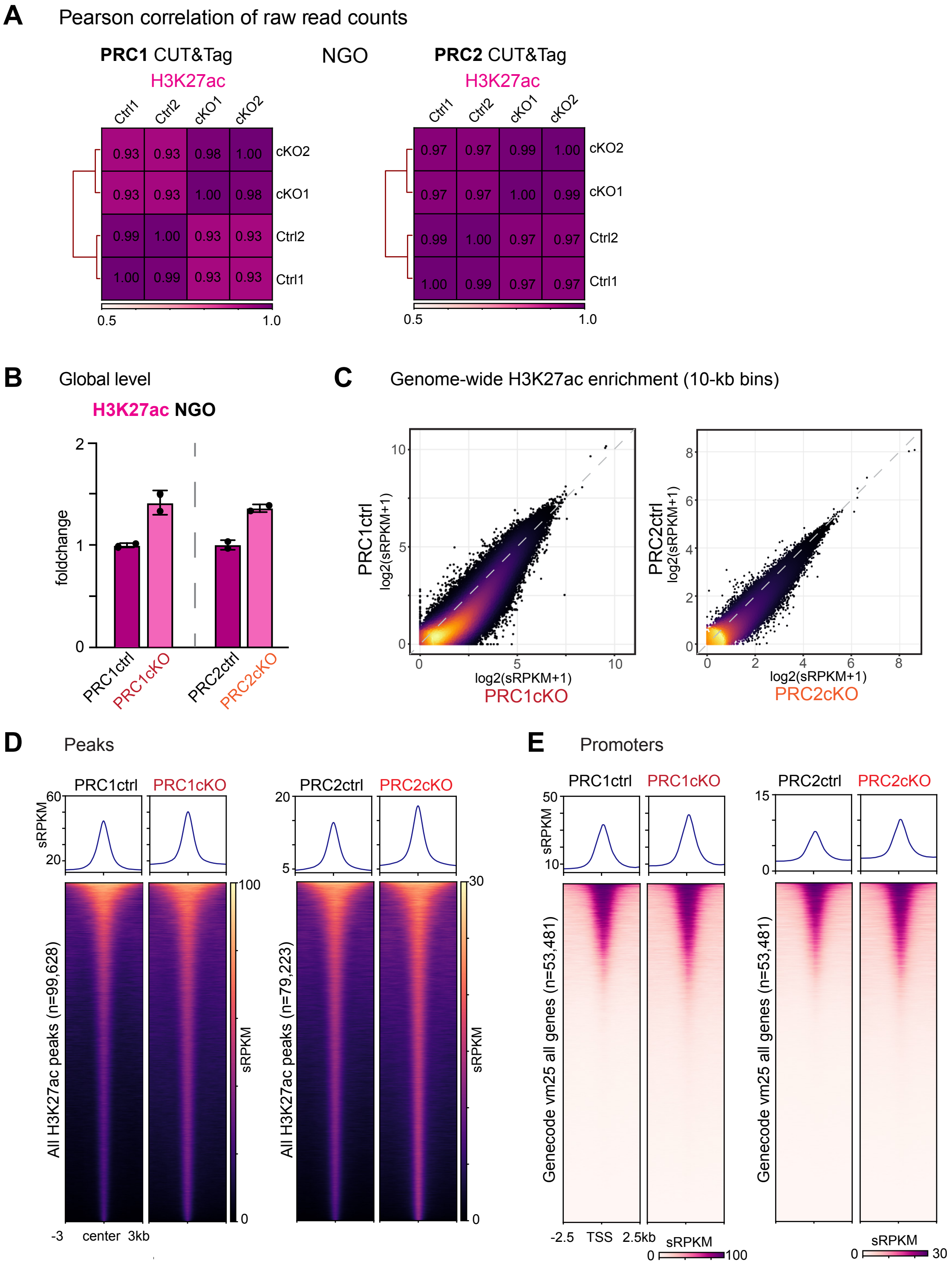
A
Pearson correlation of raw read counts
PRC1 CUT&Tag
H3K27ac
NGO
PRC2 CUT&Tag
H3K27ac
Ctrl1
Ctrl2
cKO1
cKO2
0.5
1.0
B
Global level
H3K27ac NGO
foldchange
PRC1ctrl
PRC1cKO
PRC2ctrl
PRC2cKO
C
Genome-wide H3K27ac enrichment (10-kb bins)
PRC1ctrl
log2(sRPKM+1)
PRC1cKO
PRC2ctrl
PRC2cKO
D
Peaks
sRPKM
All H3K27ac peaks (n=99,628)
All H3K27ac peaks (n=79,223)
-3
center
3kb
E
Promoters
Genecode vm25 all genes (n=53,481)
-2.5
TSS
2.5kb

**Fig. S5. H3K27ac changes in the Polycomb conditional knockout mouse models.**
(A) Heatmaps with hierarchical clustering showing the Pearson correlation of the raw read counts among each biological replicate for H3K27ac in CUT&Tag datasets of PRC1ctrl&cKO and PRC2ctrl&cKO, respectively.
(B) Bar chart showing the foldchange of H3K27ac global level in PRC1ctrl&cKO and PRC2ctrl&cKO NGO (n = 2 biological replicates, indicated by dots). Data are represented as mean ± SD.
(C) Density scatter plots showing genome-wide H3K27ac enrichment by 10-kb bins, comparing PRC1ctrl&cKO, PRC2ctrl&cKO,respectively.
(D) Average tag density plots and heatmaps showing all H3K27ac peaks in PRC1ctrl&cKO and PRC2ctrl&cKO NGO.
(E) Average tag density plots and heatmaps showing H3K27ac enrichment on all promoters in PRC1ctrl&cKO and PRC2ctrl&cKO NGO.

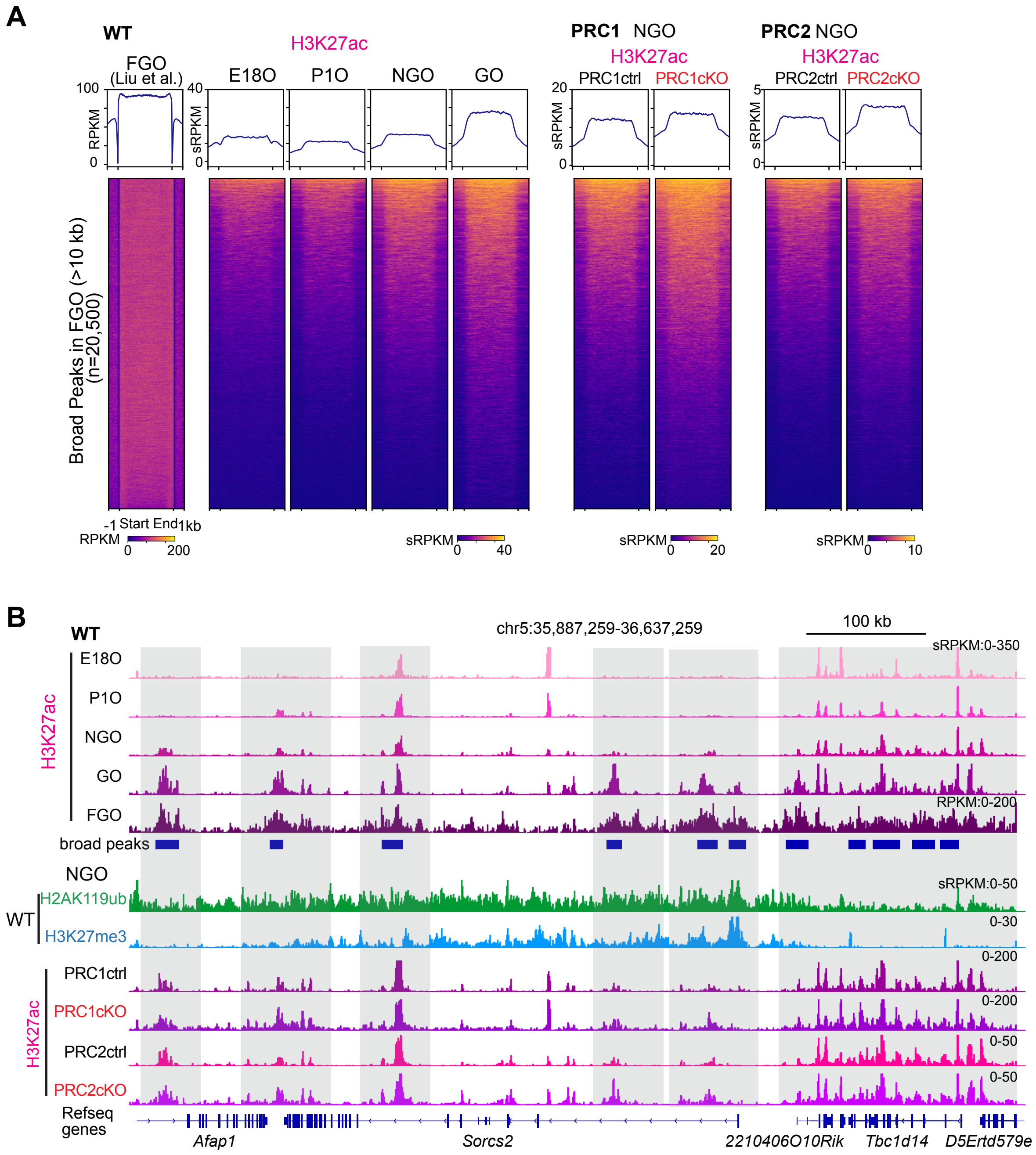


**Fig. S6. Polycomb restrains premature expansion of H3K27ac broad domains.**
(A) Average tag density plots and heatmaps showing H3K27ac dynamics in WT perinatal oocytes, PRC1ctrl&cKO and PRC2ctrl&cKO NGOs on the H3K27ac broad peaks (peaks length >10kb) identified in FGOs. H3K27ac data of WT FGOs were downloaded from GSE217970 (ref.[18]).
(B) Genome browser views of H3K27ac landscapes in different oocyte development stages of WT, and in PRC1ctrl&cKO, PRC2ctrl&cKO NGOs. Gray boxes highlight enhanced H3K27ac broad domains in PRC1cKO and PRC2cKO NGOs.

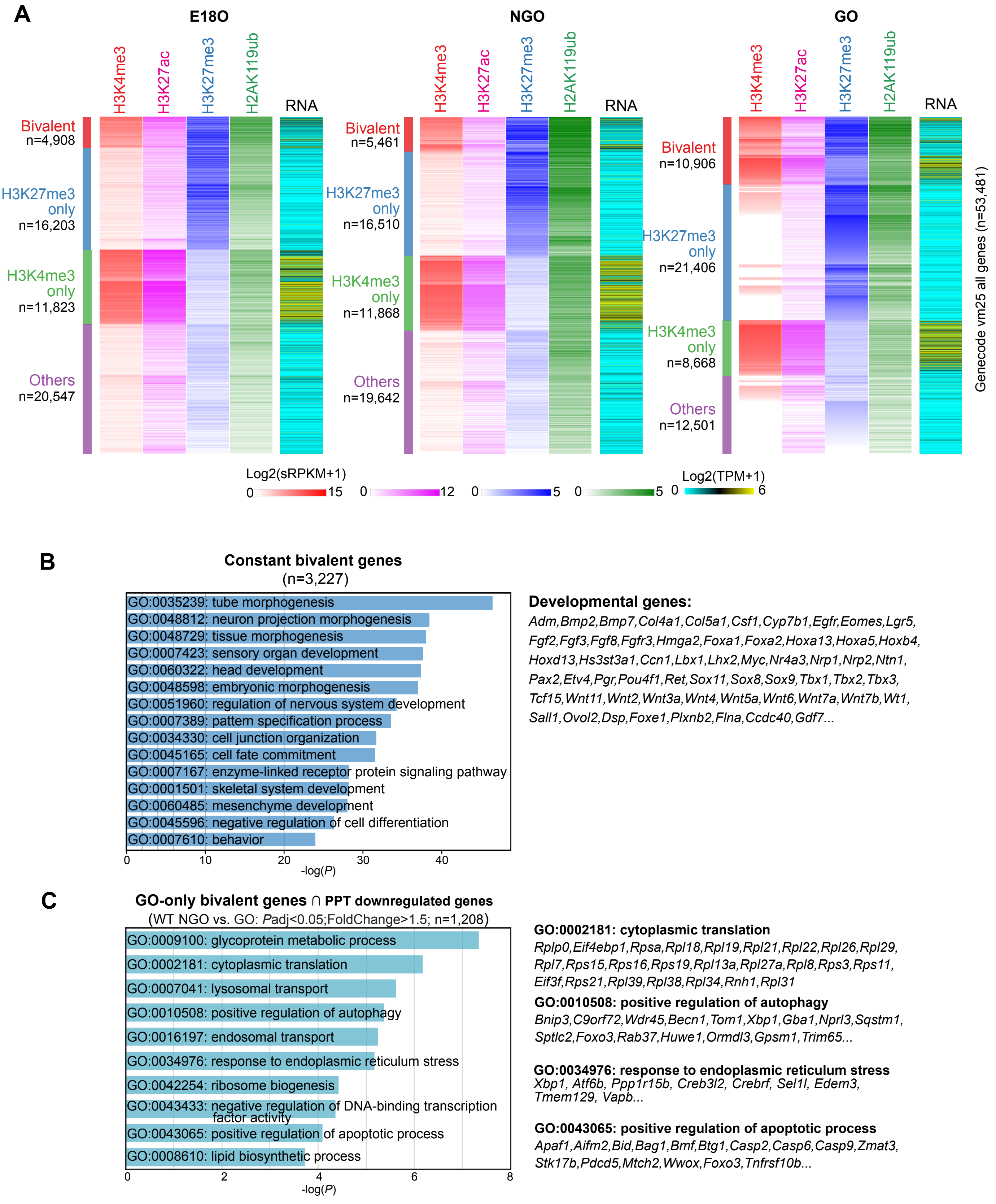
A
E18O
NGO
GO
H3K4me3
H3K27ac
H3K27me3
H2AK119ub
RNA
Bivalent n=4,908
H3K27me3 only n=16,203
H3K4me3 only n=11,823
Others n=20,547
Bivalent n=5,461
H3K27me3 only n=16,510
H3K4me3 only n=11,868
Others n=19,642
Bivalent n=10,906
H3K27me3 only n=21,406
H3K4me3 only n=8,668
Others n=12,501
Genecode vm25 all genes (n=53,481)
Log2(sRPKM+1)
0 15
0 12
0 5
0 5
Log2(TPM+1)
0 6
B
Constant bivalent genes
(n=3,227)
GO:0035239: tube morphogenesis
GO:0048812: neuron projection morphogenesis
GO:0048729: tissue morphogenesis
GO:0007423: sensory organ development
GO:0060322: head development
GO:0048598: embryonic morphogenesis
GO:0051960: regulation of nervous system development
GO:0007389: pattern specification process
GO:0034330: cell junction organization
GO:0045165: cell fate commitment
GO:0007167: enzyme-linked receptor protein signaling pathway
GO:0001501: skeletal system development
GO:0060485: mesenchyme development
GO:0045596: negative regulation of cell differentiation
GO:0007610: behavior
0 10 20 30 40
-log(P)
Developmental genes:
Adm,Bmp2,Bmp7,Col4a1,Col5a1,Csf1,Cyp7b1,Egfr,Eomes,Lgr5,
Fgf2,Fgf3,Fgf8,Fgfr3,Hmga2,Foxa1,Foxa2,Hoxa13,Hoxa5,Hoxb4,
Hoxd13,Hs3st3a1,Ccn1,Lbx1,Lhx2,Myc,Nr4a3,Nrp1,Nrp2,Ntn1,
Pax2,Etv4,Pgr,Pou4f1,Ret,Sox11,Sox8,Sox9,Tbx1,Tbx2,Tbx3,
Tcf15,Wnt11,Wnt2,Wnt3a,Wnt4,Wnt5a,Wnt6,Wnt7a,Wnt7b,Wt1,
Sall1,Ovol2,Dsp,Foxe1,Plxnb2,Flna,Ccdc40,Gdf7...
C
GO-only bivalent genes ∩ PPT downregulated genes
(WT NGO vs. GO: Padj<0.05;FoldChange>1.5; n=1,208)
GO:0009100: glycoprotein metabolic process
GO:0002181: cytoplasmic translation
GO:0007041: lysosomal transport
GO:0010508: positive regulation of autophagy
GO:0016197: endosomal transport
GO:0034976: response to endoplasmic reticulum stress
GO:0042254: ribosome biogenesis
GO:0043433: negative regulation of DNA-binding transcription factor activity
GO:0043065: positive regulation of apoptotic process
GO:0008610: lipid biosynthetic process
0 2 4 6 8
-log(P)
GO:0002181: cytoplasmic translation
Rplp0,Eif4ebp1,Rpsa,Rpl18,Rpl19,Rpl21,Rpl22,Rpl26,Rpl29,
Rpl7,Rps15,Rps16,Rps19,Rpl13a,Rpl27a,Rpl8,Rps3,Rps11,
Eif3f,Rps21,Rpl39,Rpl38,Rpl34,Rnh1,Rpl31
GO:0010508: positive regulation of autophagy
Bnip3,C9orf72,Wdr45,Becn1,Tom1,Xbp1,Gba1,Nprl3,Sqstm1,
Sptlc2,Foxo3,Rab37,Huwe1,Ormdl3,Gpsm1,Trim65...
GO:0034976: response to endoplasmic reticulum stress
Xbp1, Atf6b, Ppp1r15b, Creb3l2, Crebrf, Sel1l, Edem3,
Tmem129, Vapb...
GO:0043065: positive regulation of apoptotic process
Apaf1,Aifm2,Bid,Bag1,Bmf,Btg1,Casp2,Casp6,Casp9,Zmat3,
Stk17b,Pdcd5,Mtch2,Wwox,Foxo3,Tnfrsf10b...

**Fig. S7. Chromatin dynamics and functional annotation of bivalent promoters during perinatal oogenesis.**

(**A**) Heatmaps showing H3K4me3, H3K27ac , H3K27me3, and H2AK119ub dynamics at promoter regions (TSS ± 1 kb) and RNA expression of corresponding genes in perinatal oocytes. The color keys represent signal intensity, log2-transformed sRPKM values or TPM values.

(**B, C**) Gene Ontology analyses of the bivalent genes of indicated groups during perinatal oogenesis. Key Gene Ontology terms and representative genes are shown. *P* values were generated by Metascape using the two-sided hypergeometric test.

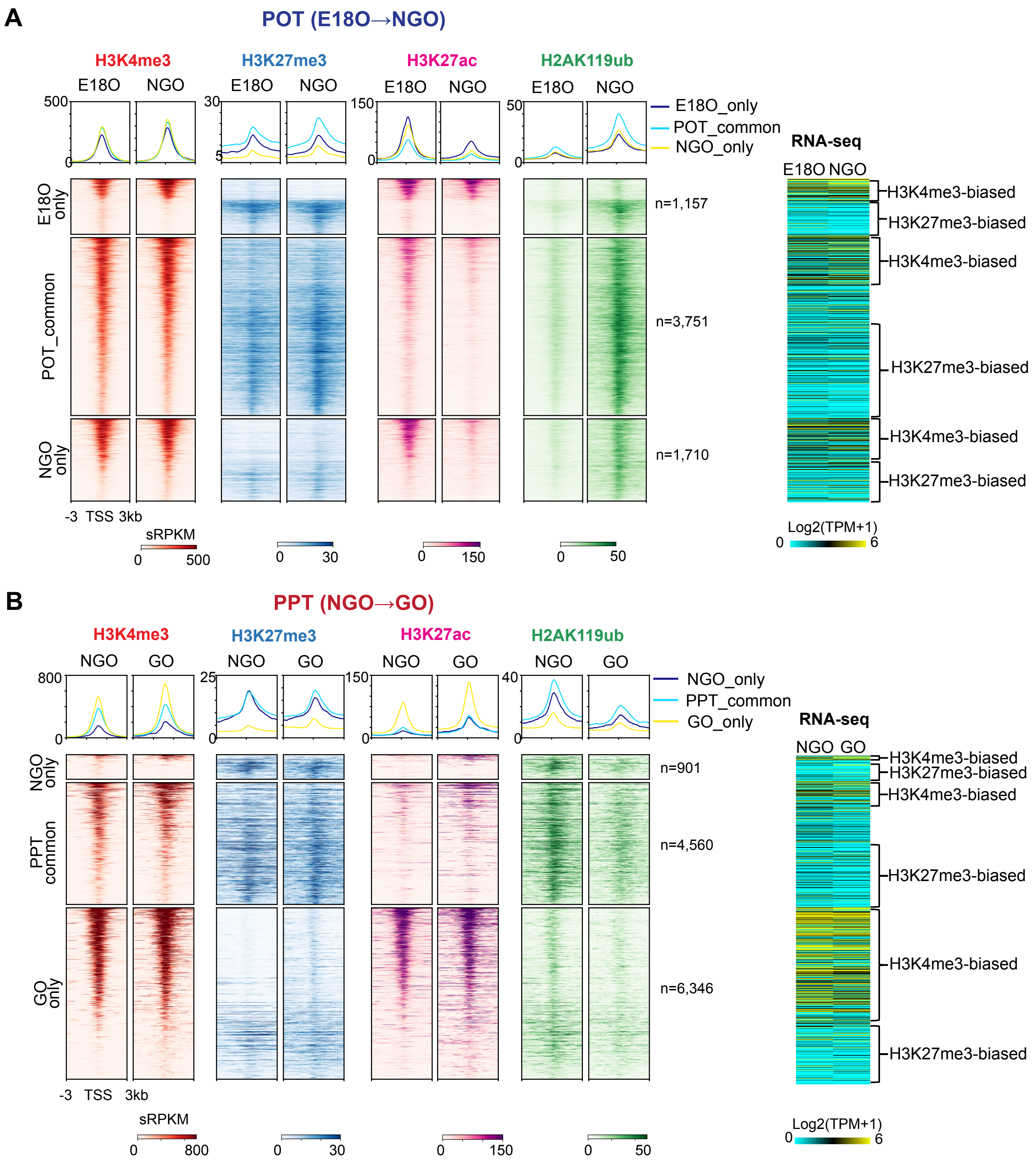


**Fig. S8. Dynamics of promoter bivalancy during POT and PPT.**
(A and B) Average tag density plots and heatmaps showing histone modification changes at bivalent promoter regions (TSS ± 3 kb) during POT (A) and PPT (B). The color keys represent signal intensity, and the numbers represent sRPKM values. Right: Heatmaps showing RNA expression (log2-transformed TPM) of corresponding bivalent genes in WT oocytes.

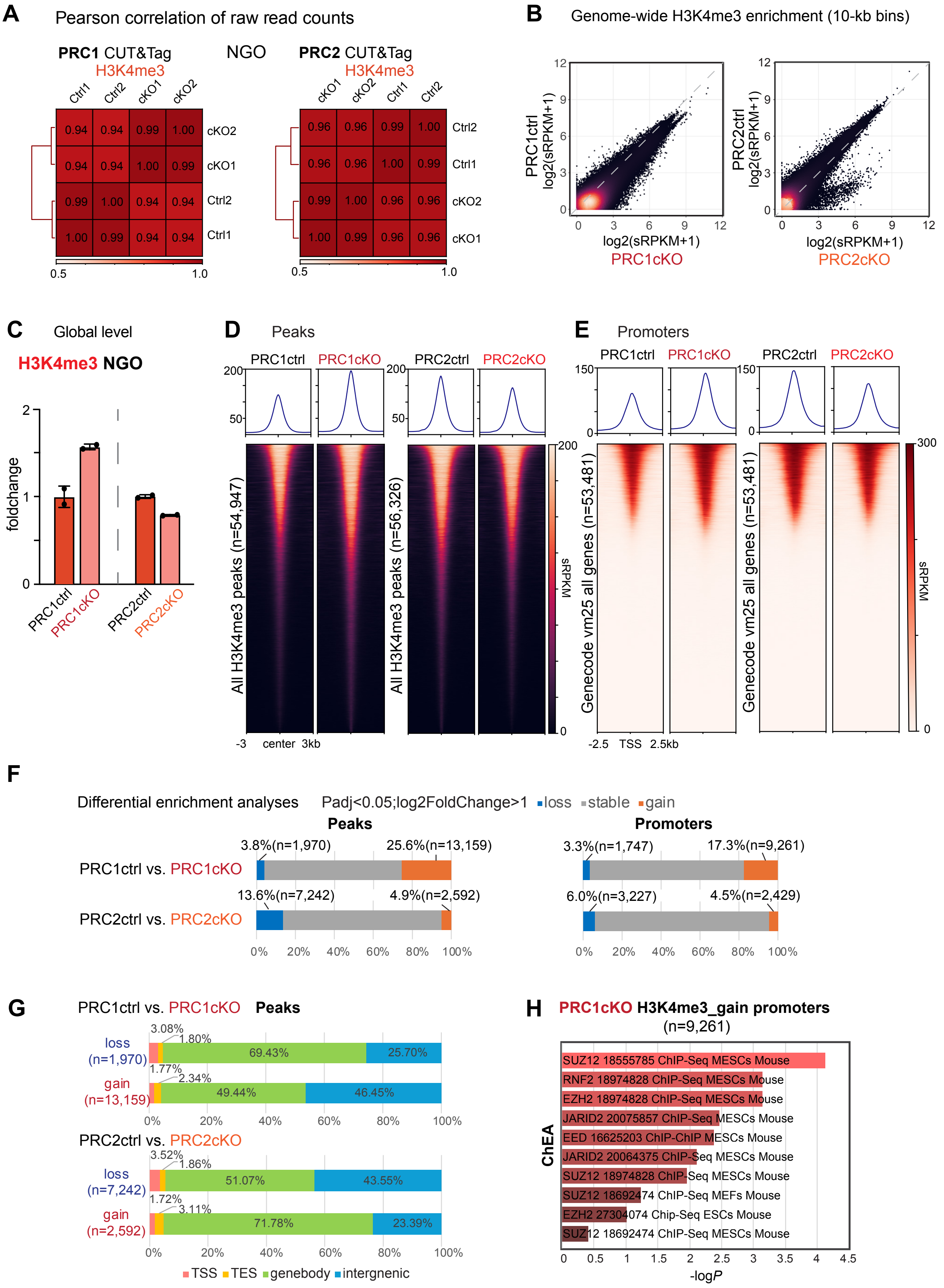

A
Pearson correlation of raw read counts
PRC1 CUT&Tag
H3K4me3
NGO
PRC2 CUT&Tag
H3K4me3
B
Genome-wide H3K4me3 enrichment (10-kb bins)
PRC1ctrl log2(sRPKM+1)
PRC2ctrl log2(sRPKM+1)
log2(sRPKM+1)
PRC1cKO
PRC2cKO
C
Global level
H3K4me3 NGO
foldchange
PRC1ctrl
PRC1cKO
PRC2ctrl
PRC2cKO
D
Peaks
All H3K4me3 peaks (n=54,947)
All H3K4me3 peaks (n=56,326)
-3 center 3kb
sRPKM
E
Promoters
Genecode vm25 all genes (n=53,481)
-2.5 TSS 2.5kb
F
Differential enrichment analyses
Padj<0.05;log2FoldChange>1
loss
stable
gain
Peaks
Promoters
PRC1ctrl vs. PRC1cKO
PRC2ctrl vs. PRC2cKO
3.8%(n=1,970)
25.6%(n=13,159)
13.6%(n=7,242)
4.9%(n=2,592)
3.3%(n=1,747)
17.3%(n=9,261)
6.0%(n=3,227)
4.5%(n=2,429)
G
PRC1ctrl vs. PRC1cKO Peaks
loss (n=1,970)
gain (n=13,159)
PRC2ctrl vs. PRC2cKO
loss (n=7,242)
gain (n=2,592)
TSS
TES
genebody
intergnenic
H
PRC1cKO H3K4me3_gain promoters (n=9,261)
ChEA
SUZ12 18555785 ChIP-Seq MESCs Mouse
RNF2 18974828 ChIP-Seq MESCs Mouse
EZH2 18974828 ChIP-Seq MESCs Mouse
JARID2 20075857 ChIP-Seq MESCs Mouse
EED 16625203 ChIP-ChIP MESCs Mouse
JARID2 20064375 ChIP-Seq MESCs Mouse
SUZ12 18974828 ChIP-Seq MESCs Mouse
SUZ12 18692474 ChIP-Seq MEFs Mouse
EZH2 27304074 Chip-Seq ESCs Mouse
SUZ12 18692474 ChIP-Seq MESCs Mouse
-logP

**Fig. S9. H3K4me3 changes in the Polycomb conditional knockout mouse models.**
(A) Heatmaps with hierarchical clustering showing the Pearson correlation of the raw read counts among each biological replicate for H3K4me3 in CUT&Tag datasets of PRC1ctrl&cKO and PRC2ctrl&cKO, respectively.
(B) Density scatter plots showing genome-wide H3K4me3 enrichment by 10-kb bins, comparing PRC1ctrl&cKO and PRC2ctrl&cKO.
(C) Bar chart showing the foldchange of H3K4me3 global level in NGO of PRC1ctrl&cKO and PRC2ctrl&cKO (n = 2 biological replicates, indicated by dots). Data are represented as mean ± SD.
(D) Average tag density plots and heatmaps showing all H3K4me3 peaks in PRC1ctrl&cKO and PRC2ctrl&cKO NGO.
(E) Average tag density plots and heatmaps showing H3K4me3 enrichment on all promoters in PRC1ctrl&cKO and PRC2ctrl&cKO NGO.
(F) Bar charts showing percentages of differentially enriched peaks, and promoters for H3K4me3 in NGO of PRC1ctrl&cKO and PRC2ctrl&cKO.
(G) Genomic distribution of diffenrential enriched H3K4me3 peaks in NGO of PRC1ctrl&cKO and PRC2ctrl&cKO.
(H) Bar charts of ChIP-x Enrichment Analysis (ChEA) of gene promoters gaining H3K4me3 differentially in PRC1cKO NGO. ChEA enrichment shows the transcription factors and the cell and animal types used in the profiling experiments.
